# Same Sentences, Different Grammars, Different Brain Responses: An MEG study on Case and Agreement Encoding in Hindi and Nepali Split-Ergative Structures

**DOI:** 10.1101/2024.02.12.579942

**Authors:** Dustin A. Chacón, Subhekshya Shrestha, Brian W. Dillon, Rajesh Bhatt, Diogo Almeida, Alec Marantz

## Abstract

At first glance, the brain’s language network appears to be universal, but languages clearly differ. How does the brain adapt to the specific details of individual grammatical systems? Here, we present an MEG study on case and agreement in Hindi and Nepali. Both languages use split-ergative case systems. However, these systems interact with verb agreement differently – in Hindi, case features conspire to determine which noun phrase (NP) the verb agrees with (subject, object, or neither), but in Nepali the verb always agrees with the subject NP. We found that left inferior frontal and left anterior temporal regions are sensitive to case features in both languages. Across case configurations, these same brain areas in Hindi participants show different patterns of activity for sentences that require masculine vs. feminine marking on the verb, before the comprehenders encounter it. Additionally, the left temporoparietal junction in Hindi shows different activity for subject and object agreement configurations. Both findings are not observed in Nepali participants. We suggest that this brain response demonstrates a unique-to-Hindi selection of an agreement controller and pre-encoding of the verb’s morphological features. This shows that brain activity reflects psycholinguistic processes that are intimately tied to grammatical features.

**Highlights:**

- The left inferior frontal lobe and the left anterior temporal lobe distinguish accusative objects versus bare object NPs in Hindi and Nepali, and pre-emptively encode gender agreement features in Hindi.
- The left inferior parietal lobe shows a differential sensitivity to object-agreement and subject-agreement constructions in Hindi that is absent in Nepali
- MEG can reveal differences in neural activity that reflect specific requirements of different grammatical systems

## 1. Introduction

Languages can appear quite different from one another. Nonetheless, a typical human can learn and use any natural human language. The organization of the brain’s “language network” appears to be largely uniform across languages, despite the diversity in features they may have, or the kinds of information that their structures encode (Dunagan et al. 2022; Malik-Moraleda et al. 2022), although some results suggest some differences (Wei et al. 2023). Given the reasonable assumption that neural structures and language processing strategies are essentially the same across languages, differences in neural activity should reflect the way that a uniform representational and processing system deals with parametric differences between grammatical systems. For example, a predictive processing mechanism will predict the arguments of a verb from the verb in a subject-verb-object (SVO) language, whereas the verb might be predicted from both arguments in an subject-object-verb (SOV) language, or languages with strict word order and no case marking may weight word order as a stronger predictive cue than languages with freer word order and case marking (e.g., Schlesewsky & Bornkessel-Schlesewsky 2009). Identifying systematic differences in neural activity in language processing between languages can serve as a key strategy for relating grammatical descriptions, processing mechanisms, and neurobiology.

We examine this question through the lens of case and agreement dependencies. Case morphology, such as the difference between nominative *he* and accusative *him*, provide the comprehender with important cues to ‘who-did-what-to-whom’ via syntactic relations. Like case, agreement also guides the comprehender in identifying entities in the sentence. In English, the verb agrees (corresponds in form) with the subject noun phrase (NP) in number – *the boy quickly run-**s**, the boy-**s** quickly run*. However, different languages require that the verb relates to its arguments differently. For instance, in many languages, such as Hindi, Nepali, Arabic, or Hebrew, the verb agrees with its subject in gender, unlike English – *the boy quickly run-**s**, the girl quickly run-**s***.

Here, we leverage the “split-ergative” agreement systems of two very typologically similar languages: Hindi and Nepali. Hindi and Nepali are both predominantly SOV languages from the Indo-Aryan language family. In both languages, the subject noun phrase (NP) requires an ergative case suffix (ने *-ne* in Hindi, ले *-le* in Nepali) in simple past (perfective) sentences, but not in the simple (imperfective) present, imperfective past, or future. In the simple present, the verb agrees with the nominative subject in number, gender, and person, much like Western European languages. However, in the perfective past, the languages use different rules. In Hindi, the verb usually agrees with the object in the simple past, and not with the subject carrying the ने*-ne* suffix. In Nepali, the verb also agrees with the subject carrying the ले *-le* suffix in the past tense, i.e., no difference is observed between present and past tense in Nepali.

These grammatical features mean that different information is accessible at different points in Hindi and Nepali, in sentences that are otherwise structurally similar. In Nepali, comprehenders can identify the expected number, gender, and person of the verb shortly after encountering the subject NP, regardless of its case. However, in Hindi, the comprehender’s expectations of the verb’s form differs depending on the presence of the ergative suffix ने *-ne.* If the subject NP is bare, then Hindi comprehenders know that the verb will match with the subject NP in number, person, and gender, just as in Nepali. If the subject NP has the ergative suffix ने *-ne,* then the comprehender must wait to encounter the object NP to determine the verb’s features. In other words, Nepali comprehenders can always identify the subject NP as the ‘agreement controller’, i.e., the noun with which the verb must match in features. By contrast, Hindi comprehenders must examine the morphology of the subject NP and object NP to determine which NP is the controller, and may not know the likely agreement specification on the verb until after integrating both NPs into the parse of the sentence.

Here, we investigate the relation between brain activity and the different accessibility of agreement-relevant cues during sentences processing in Hindi and Nepali using magnetoencephalography (MEG). We focus on the evoked brain activity during processing of the object NP, prior to the verb, in simple SOV sentences in ergative-perfective structures and nominative-imperfective structures, focusing on NP-verb agreement in gender marking. We assume that Hindi comprehenders explicitly identify and store in memory an argument NP as the agreement controller (Bhatia 2019; Bhatia & Dillon 2022), which is not coextensive with a specific grammatical role (subject, object) or case morphosyntax (e.g., nominative). This is consistent with the view that agreement processing depends on anticipatorily encoding features of the verb, including agreement features such as gender, prior to encountering it (Eberhard et al. 2005; Chacón 2022; Keshev & Meltzer-Asscher 2024). Due to the agreement rules of Hindi, the expected features of the verb cannot be uniformly identified until the object NP, i.e., after the case features of both principal arguments have been identified and integrated into a syntactic structure. By contrast, Nepali comprehenders should be able to identify the likely morphosyntactic features on the verb shortly after processing the subject NP, and thus these predictive processes do not need to be present during the processing of the object NP.

The neuroanatomical correlates of the processes we investigate here are not well-understood. We have two key hypotheses. The first hypothesis is that subject and object case morphology will interact in the brains of Hindi comprehenders during the processing of an object NP. Specifically, we expect that the processing of a bare object NP that controls agreement vs. the processing of a bare object NP that does not will show different patterns of activity. We do not have *a priori* hypotheses about the directionality of this effect, nor the region that this will occur in, although we suspected that it would be in a key area associated with case-agreement processing: left prefrontal regions, left anterior temporal regions, and inferior parietal regions. If this interaction in case features corresponds to identifying an agreement controller, according to the split-ergative agreement system observed in Hindi, then this interaction of case features should not be observed in Nepali. The second hypothesis is that Hindi comprehenders may not encode the expected inflectional features of the verb until the object NP has been processed, at least in sentences with ergative subject NPs. Thus, we expect divergent MEG signals between sentences that require a masculine vs. feminine verb after the object NP, orthogonal to the gender specifications of any particular argument NP across sentence types. Here too we had no specific prediction about the directionality of the effect or the localization in neuroanatomy, although we again suspected that this would likely occur in one of the key regions associated with case-agreement. We presume that Nepali comprehenders also formulate an expectation for the agreement specification on the verb, but we did not predict that this needs to be observable during the processing of the object NP. Moreover, because the verb agrees with the subject NP regardless of whether its case is nominative or ergative, the patterns of brain activity would be underdetermined with respect to indexing the expected gender features of the verb or the gender features of the subject NP. Crucially, we make no *a priori* hypotheses about main effects of ergative vs. nominative case during the processing of the subject NP or accusative vs. bare case during the processing of the object NP. There are likely many differences engendered by these features in these time windows, and instead we focus our hypothesis on the interaction of these features during the processing of the object NP period. AA summary of our findings are presented in Figure 1.

**Figure 1.**
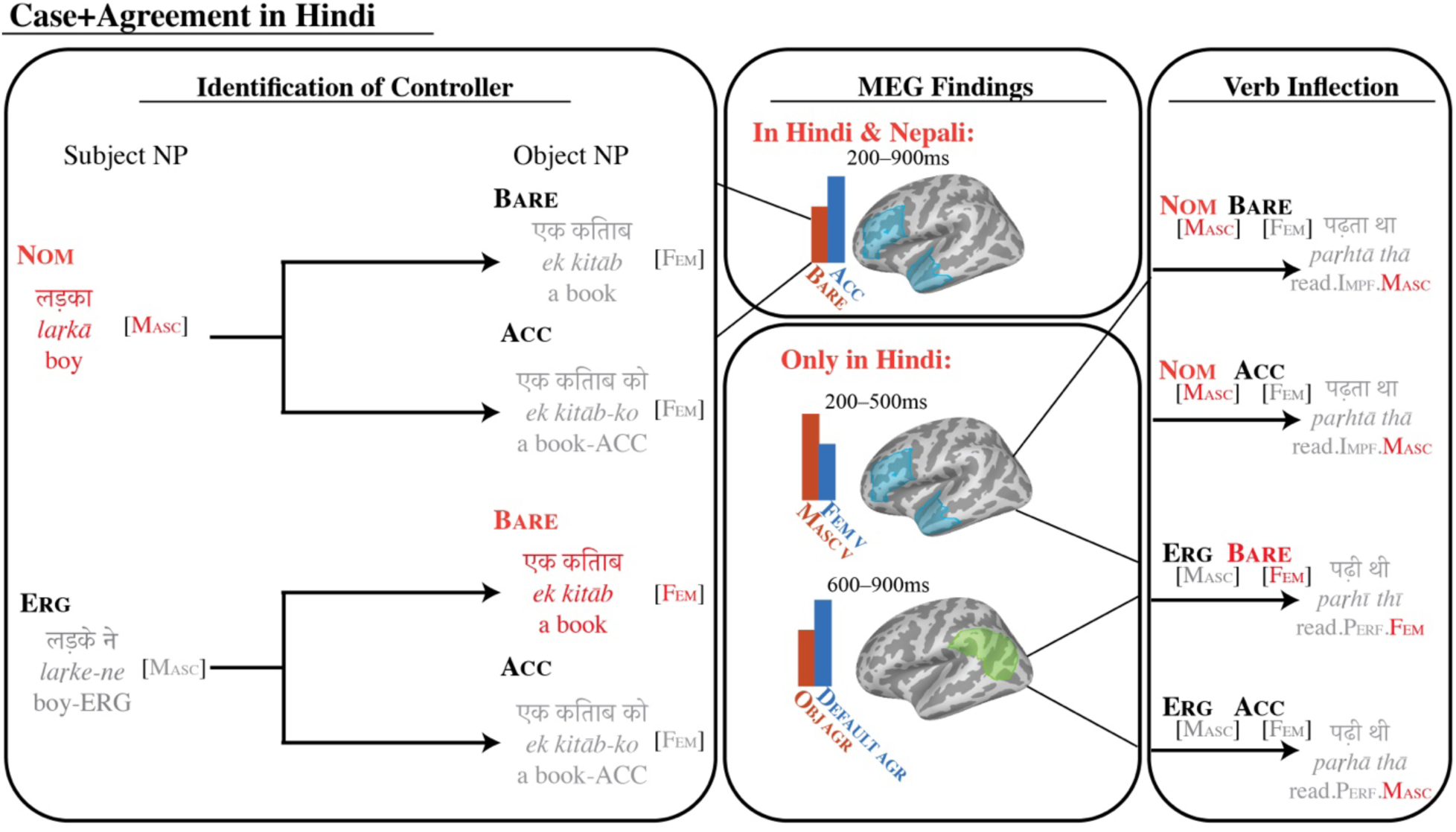
Summary of key findings. During the processing of an object NP in Hindi, left anterior temporal and left inferior frontal regions show greater activity for accusative object NPs compared to bare object NPs, 200–900ms. Similar findings are observed in Nepali. Neural activity in these same regions diverge for sentences that will result in a masculine vs. feminine verb, 200–500ms in Hindi. Object- and default-agreement configurations show distinct patterns of activity in left inferior parietal lobe, 600–900ms.

### 1.1 Case-and-Agreement Processing and Predictive Encoding

Subject-verb agreement dependencies are extensively studied in psycholinguistics as a window into the memory operations that support language comprehension generally (Nicol et al. 1997; Badecker & Kumiak 2007; Wagers et al. 2009; Dillon et al. 2013), and as a window into the kinds of representations that are built in real-time (Bock & Miller 1991; Franck et al. 2002; Bhatia & Dillon 2022; Chacón 2022). Subject-verb agreement processing likely consists of at least two distinct stages; the first involves identifying possible agreement controllers and predicting likely morphosyntactic features of the upcoming verb (Eberhard et al. 2005; Wagers et al. 2009; Bhatia & Dillon 2022; Chacón 2022), while the second involves retrieval of the subject NP’s features after the verb is encountered (Nicol et al. 1997; Bedecker & Kumiak 2007; Wagers et al. 2009; Dillon et al. 2013).

Electroencephalography (EEG) recordings demonstrate rapid detection of verbs that satisfy vs. violate agreement rules (Hagoort et al. 1993; Friederici 2002; Nevins et al. 2007; Choudhary et al. 2009; Molinaro et al. 2011; Bhattamishra et al. 2021). Agreement violations, in which the verb and its argument mismatch in a feature, usually elicit a biphasic response, with a left anterior negativity (LAN) ∼300ms post-verb onset, followed by a P600, a positive deflection ∼600ms post-verb onset. These earlier responses may reflect rapid detection of a violated predicted verb form (Molinaro et al. 2011). Similarly, using MEG, Tucker et al. (2014) found greater activity for verbs that satisfy agreement rules vs. ungrammatical verbs ithat mismatched the subject n number in the left posterior superior temporal lobe (lpSTL), ∼700ms post-verb onset.

Hemodynamic measures investigating agreement violations have implicated a variety of brain regions in agreement configurations, including the LIFL, portions of the left anterior temporal lobe (LATL) and left posterior temporal lobe (LPTL), and LIPL and its right hemisphere homologs. In an fMRI study investigating case and agreement violations in Basque, a language with ergative case and both obligatory subject-verb and object-verb agreement, Nieuwland et al. (2011) observed greater activation in the LIPL and its right hemisphere homologs (RIPL) for object-verb agreement violations. They also found greater activation for object-verb mismatch in the left and right middle frontal lobes (LMFL, RMFL). In an fMRI study in Spanish, a language with obligatory gender- and number-agreement between nouns and modifiers, Carreiras et al. (2015) found greater activity in LIFL for adjective-noun and determiner-noun pairs that mismatched in gender or number. They additionally found greater activation in right parietal areas for determiner- and adjective-noun pairs that mismatched in number (see also Carreiras et al. 2010b). Similarly, in an fMRI study in Spanish, Quiñones et al. (2014) identified greater activity in LIPL and LMFL for ungrammatical subject NP-verb agreement. They also identified greater activity in LATL, LpSTL, LIFL, and portions of the left orbitofrontal cortex (OF) for grammatical structures that exceptionally allow for a mismatch in person features (‘unagreement’). Quiñones et al. (2018) found that grammatical nouns with transparent gender marking (*libr-**o*** ‘book’; *-o* is the masculine gender marker) engaged LPTL and LIPL and the right homologs (RPTL, RIPL) more than ‘opaque’ nouns (*lápiz* ‘pencil’), but opaque nouns resulted in greater activity in bilateral frontal and superior temporal lobe. Finally, in an fMRI study in Japanese, Cui et al. (2022) found different patterns of activity for verb marking corresponding to the relative social status of the speaker vs. other NP referents in left frontal regions, LATL, and LIPL.

In direct neuron recordings, Sahin et al. (2009) found greater activity in the LIFL when participants produced English inflected verb and noun forms ∼300ms after word-onset (*rock-s, rock-ed*) compared to uninflected forms. Similar findings were observed using production study using fMRI by Kielar et al. (2011). Using a similar paradigm in MEG with word production in English, Hauptman et al. (2021) observed greater activity in LIFL and LATL corresponding to verb and noun inflection, approximately 300ms after the cue to begin speaking was provided.

However, due to the limitations of these methodologies, it has been difficult to draw firm inferences about which brain areas are responsible for which computations. EEG recordings have poor spatial resolution, meaning it is difficult to identify the neural generators for the measured activity. Inversely, hemodynamic measures afford poorer temporal resolution, rendering it difficult to disentangle memory retrieval operations encountered at the verb versus pre-verbal computations supporting selection of agreement controllers and pre-emptive encoding of verb features. Finally, although direct neuron recordings and MEG recordings enjoy high spatial and temporal resolution, these studies have predominantly focused on computations executed at the verb, rather than anticipatory processes that might occur before the verb is encountered, e.g. identifying a controller and predicting the verb’s features. Thus, little is understood about how the brain executes these computations. Nonetheless, these studies demonstrate that LIFL, LATL, and LIPL are likely candidates in processing argument-verb agreement relations.

The hypothesis that we entertain in this paper is that comprehenders of Hindi exploit morphosyntactic case-marking as a cue for predicting agreement features, and that these processes are different in Nepali. Overwhelmingly, comprehenders appear capable of integrating many cues to predict upcoming semantic and syntactic features of an upcoming verb (cf. Ferreira & Qiu 2021). In EEG recordings, greater negative amplitude around 200–400ms, the ‘N400’, is associated with less predictability of a word given its prior the context (*I drink my coffee with sugar and socks / cream*). The MEG correlate of the N400, the ‘M350’, and fMRI responses to unexpected words localize the effect of responses to unpredictable words in the left middle temporal lobe (LMTL), although left frontal and LOF regions also are crucial for detecting unpredicted words (Baumgaertner et al. 2002; Halgren et al. 2002; Pylkkänen et al. 2004; Van Petten & Luka 2006; Lau et al. 2008; Lau & Namyst 2019).

Given that comprehenders are capable of integrating many cues into their predictions, then comprehenders of different languages should attend to different cues, depending on the structural features of their languages. This has been demonstrated robustly over a series of studies by Bornkessel-Schlesewsky and colleagues (Bornkessel-Schlesewsky & Schlesewsky 2008, 2009; Bornkessel-Schlesewsky et al. 2016). For instance, they argue that users of German, which exhibits flexible word order and case-marking, rely on case information and animacy more than word order to guide their interpretation of NPs (Schlesewsky & Bornkessel-Schlesewsky 2009). In an fMRI study in German, a language with freer word order and case marking, Bornkessel et al. (2005) found greater activity for sentences with unlikely word order and verb combinations compared to more likely combinations in left frontal regions, lpSTL, and left ventral premotor cortex. Additionally, they found that German NPs with ambiguous case (*Lehrerinnen* ‘teachers’; may be dative or nominative) modulate neural activity differently in LIPL and lPSTL than NPs with unambiguous case marking (*den*[DAT] *Lehrerinnen*[DAT] ‘(to) the teachers’). Mandarin Chinese, by contrast, is a language that overwhelmingly prefers SVO word order, and features no case or agreement morphology. Mandarin Chinese users therefore heavily rely on word order and animacy as cues to interpreting sentences (Wang et al. 2009; Wang et al. 2012), although this too can be modulated by the demands of grammatical constructions (e.g., ‘adversative passives’ in Philipp et al. 2008). Similarly, Gattei et al. (2015) found that in Spanish, a language with flexible word order like German but with less explicit case morphology, comprehenders user both word order and case marking to infer the likely semantic class of the verb in ‘marked’ word order-case configurations that are highly associated with particular verbal semantics (see also Simonsen & Chacón 2024).

The majority of work on prediction in language comprehension contrasts the neural responses to predicted vs. unpredicted material. What do we know about the mechanisms of predictions themselves? In an MEG study on two-word adjective-noun phrases in English with highly predictive adjectives (e.g., *stainless* strongly predicts *steel*), Fruchter et al. (2015) found that the predictiveness of adjectives (e.g., *p*( ‘steel’ | ‘stainless’)) correlated with MEG activity in LATL regions 100–500ms post-adjective onset, and LIFL regions 400–500ms post-adjective onset. Similarly, Wang and colleagues also examined the neural activity prior to a predicted word to determine the neural correlates of committing to a predicted word. They compared the neural activity in EEG and MEG recordings after highly predictive verbs vs. less predictive verbs in Mandarin Chinese (Wang et al. 2018) and English (Wang et al. 2020). Using a representational similarity analysis (RSA; Kriegeskorte et al. 2008), they found greater similarity among the spatial patterns of the evoked responses 400–600ms after highly constraining verbs compared to less constraining verbs, before the predicted noun was encountered.

Our prediction specifically concerns the early encoding or predicting of agreement features of the verb given the case-marking on the preceding NPs. But, what are the neuroanatomical correlates of the processes implicated in processing case? As mentioned earlier, fMRI studies in German and Basque point to lpSTS, LIPL, RIPL, and bilateral MFL as key regions. Additionally, in an fMRI study in English, Yokoyama et al. (2012) found greater activation for LIFL and LMFL regions, LIPL, and LMTL when participants were instructed to select the appropriate case form in a sentence context. Thus, there appears to be significant overlap between the neuroanatomical regions implicated in agreement processing (LIFL, LATL, LIPL), case processing (LIPL, LIFL), and lexical prediction (LPTL, LATL).

### 1.2 Split-Ergativity in Hindi and Nepali

Our study focuses on Standard Hindi and Nepali. Both languages are typically written in the Devanagari script and use SOV word order. Hindi nouns, verbs, and adjectives are marked for masculine or feminine gender (typically with *-ā* masculine singular, *-e* for masculine plural, and *-ī* for feminine). Nepali animate nouns, verbs, and adjectives are marked for gender (typically *-ā* and *-o* for masculine and *-ī* and *-i(n)* for feminine), and verbs inflect for number. Inanimate nouns typically default to masculine in Nepali.

Both languages use a split-ergative case system. In the perfective aspect, the subject NP normally must end in the ergative case suffix ने *-ne* in Hindi and ले *-le* in Nepali. In other tense/aspect combinations, the subject NP normally appears in the nominative case, which is not associated with a suffix. For concreteness, we refer to the argument of intransitive predicates and the agentive or external argument of transitive predicates as ‘the subject NP’, regardless of case and agreement, and we refer to the theme, patient, or internal argument of a transitive predicate as ‘the object NP’, although these labels can be problematic for some ergative languages (Dixon 1994; Aldridge 2008; see also Carreiras et al. 2010a and Laka 2012; for relevance to language processing with respect to Basque; Polinsky et al. 2012 for Avar, and Logenbaugh & Polinsky 2016 and Tollan et al. 2019 for Niuean). Unlike in other languages, ergative subjects in Hindi and Nepali exhibit more of the ‘typical’ properties of subjects compared to their nominative counterparts (e.g.. de Hoop & Swarts 2009 for Hindi).

Although the correlation between aspect and subject case morphology is strong, it is not absolute in either language. In Hindi, the ergative suffix is optional with some intransitive verbs in the perfective aspect (such as खाँस - *khāṃs-* ‘cough’), i.e., even if there is only one NP argument. Some so-called ‘light verbs’, verbal auxiliaries that combine with other verbs to modulate their aspectual interpretation, require the assignment of ergative case, and others exceptionally require that the subject is bare, even for transitive predicates in the perfective aspect (Kachru 1980; Mahajan 2012). Lastly, Hindi users may choose to mark the agent with ergative case to assert that the event was intentional, or may choose to omit it to express a non-volitional action in some contexts (Butt & Deo 2017). In Nepali, the ergative suffix ले *-le* may also appear with verbs not in the perfective aspect, due to an array of factors (see Li 2007 for overview). Nepali users may use the ergative suffix to disambiguate whether an inanimate or non-human NP is the agent of the action (Abadie 1974), to focus or ‘emphasize’ the subject (Bickel 2011), to modulate the relationship between the subject and predicate, i.e., to express an individual-level predicate (Butt & Poudel 2007) or to express a ‘categorical proposition’ (Kuroda 1972; Lindemann 2019). Unlike in Hindi, light verbs are reported to not affect the case morphology of the subject (Slade 2014). Despite these complications, EEG evidence from Hindi demonstrates that ergative case morphology is rapidly integrated as a predictive cue for perfective aspect. For instance, Choudhary et al. (2009) found that evoked responses at the verb exhibited increased N400 and P600 amplitudes for imperfective verbs occurring after an ergative subject, and an increased N400 amplitude for perfective verbs occurring after a nominative subject. Similarly Matthew et al. (2024) reported that subject NP case morphology is a predictive cue for transitivity and light verb choice in Hindi, given the interaction between these elements.

In addition to this subject NP case alternation, both languages also exhibit an object NP case alternation. Animate object NPs obligatorily surface with the accusative/dative case suffix को -*ko* in Hindi and लाई -*lāī* in Nepali. Inanimate object NPs may occur bare or with the accusative case. Inanimate object NPs that are marked with the accusative case carry a specific or definite interpretation. The variation in case of the subject NP and object NP are orthogonal, and do not appear to change their thematic interpretation or the grammatical relations between the NP arguments and the verb apart from agreement.

Hindi verb agreement is descriptively governed by what Bhatia (2019) calls The Hindi Agreement Generalization, based on Pandharipande & Kachru (1977): The verb agrees with the most structurally prominent NP that is unmarked for case. This results in an interaction between ergativity/aspect marking and agreement. In the imperfective aspect (e.g., simple present), the subject NP does not end in ergative ने *-ne*. The subject NP controls agreement in this configuration. In the perfective aspect, the subject must normally bear ergative case. In these contexts, the subject NP is blocked from controlling agreement. Instead, the next prominent NP must be considered. If the object NP is bare, then the object controls agreement, since it is now the most prominent NP without case. However, if the object NP is marked with accusative case को *-ko* to mark animacy or specificity, then the object NP is also ‘blocked’ from agreement. If both subject and object NPs are marked with case, then the verb must surface as a third person, singular, masculine ‘default’ form. In Nepali, by contrast, verbs always agree with subject NP, regardless of whether it is ergative or bare. More exhaustive studies on case and agreement in Hindi can be found in Butt and King (2004), Bhatt (2005), and Mahajan (2017).

Previous studies on the processing of Hindi/Urdu using EEG have revealed a similar rapid sensitivity to agreement violations as observed in other languages. In an agreement violation paradigm, Nevins et al. (2007) found that subject-verb agreement violations produced a greater P600 response compared to grammatical controls. This was observed for verbs that mismatched in number, gender, and person with the subject NP, with the greatest amplitude difference corresponding to person+gender mismatches. In a study leveraging the correspondence between subject NP case and verb tense/aspect, Choudhary et al. (2009) found distinct ERP patterns to verbs that mismatch in aspect with the expected case marker. Bare subject NPs followed by an ungrammatical simple perfective verb produced greater N400 amplitude post-verb onset compared to grammatical controls, and ergative subject NPs followed by an ungrammatical imperfective verb produced both greater N400 and P600 responses post-verb onset (see also Bickel et al. 2015). Bhattamishra et al. (2021) found an increase in P600 amplitude for gender agreement errors for animate subject NPs, but a negativity observed in posterior midline sensors for gender agreement with inanimate subject NPs (see also Gulati & Choudhary 2023 for different results in Punjabi). In a production study, Sauppe et al. (2021) found that EEG signatures diverged before the utterance of sentences beginning with ergative vs. bare subject NPs. Studies on Basque, another split-ergative language in which verbs agree with both subject and object NPs, reveal similar rapid detection of agreement violations for both subject- and object-agreement violations (Zawizseski & Friederici 2009; Díaz et al. 2016; but see Chow et al. 2019).

We follow Bhatia & Dillon’s (2022) proposal that Hindi comprehenders actively identify and encode an argument NP as the agreement controller. Upon encountering an object NP, Hindi and Nepali comprehenders must both identify the morphosyntactic form of the object NP, and integrate into a syntactic parse and semantic interpration of the sentence. As a subprocess, they must identify the morphosyntactic features of the head noun (person, number, and gender features, and case inflection), and access the head noun’s lexical semantics. We assume that these processes are not specific to either language. Brain activity during this period may reflect differences in terms of the morphological form of accusative objects vs. bare objects due to the visual and linguistic form of the object NP, its effect on the entire sentence (e.g. predictability of the verb, construction of a discourse interpration, etc.), or differences in interference between the subject and object NPs. Crucially, Hindi comprehenders must then identify the expected agreement controller of the verb according to different principles than Nepali comprehenders. In processing object agreement structures (ergative subject NP, bare object NP; NP-*ne* NP), Hindi comprehenders must recognize the object NP as the agreement controller, by encoding it in memory with a feature that may be used as a cue for memory retrieval later at the verb (i.e., [CONTROLLER]; Bhatia & Dillon 2022). Furthermore, assuming that agreement processing involves active encoding of expected verb features (Eberhard et al. 2005; Keshev & Meltzer-Asscher 2024), we hypothesize that comprehenders encode an (expected) form of the verb, including its gender, number, and inflectional features, and its aspect (Choudhary et al. 2009).

Hindi comprehenders must momentarily suppress the features of the subject NP in favor of those of the object NP in short-term memory, and must identify the upcoming verb’s morphological features with those of the object NP. This predicts that the evoked response to the verb should reflect the expected features of the upcoming verb during the processing of the object NP. Importantly, the identification of an agreement controller and commitment to the form of the verb presumably happens in Nepali, but it does not need to be time-locked to the processing of the object NP’s morphological features given the irrelevance of the object NP to agreement processing.

## 2. Methods

### 2.1 Participants

Twenty-five self-identified native users of Hindi and twenty-four self-identified native users of Nepali were recruited from the Abu Dhabi community. The Hindi-speaking participants were 19– 42 years old (Mean: = 28, SE = 1.4), and the Nepali-speaking participants were 18–42 years old (Mean = 24, SE = 1.4). All participants had normal or corrected-to-normal vision, and all participants were right-handed except for one Hindi-speaking participant. The Hindi-speaking participants were 7 females and 17 males, and the Nepali-speaking participants were 6 females and 18 males. The study was formally approved by the Institutional Review Board of NYU Abu Dhabi and all participants gave written consent. Of the Hindi native-users, 1 self-identified as a native users of Spanish, 6 of English, 1 of Pashto, 2 of Gujarati, and 2 of Nepali. Of the Nepali native-users, 3 self-identified also as native users of Hindi, and 1 of Doteli.

### 2.2 Materials

We prepared fifty sets of items in both Hindi and Nepali. Each item set consisted of eight simple transitive sentences, in SOV word order, written in the Devanagari script. We manipulated the Subject Case (Nominative/Ergative), Object Case (Bare/Accusative), and Verb Stem Cloze Probability (High/Low), yielding a 2×2×2 design, in both languages. These are exemplified in Table 1.

**Table 1.**
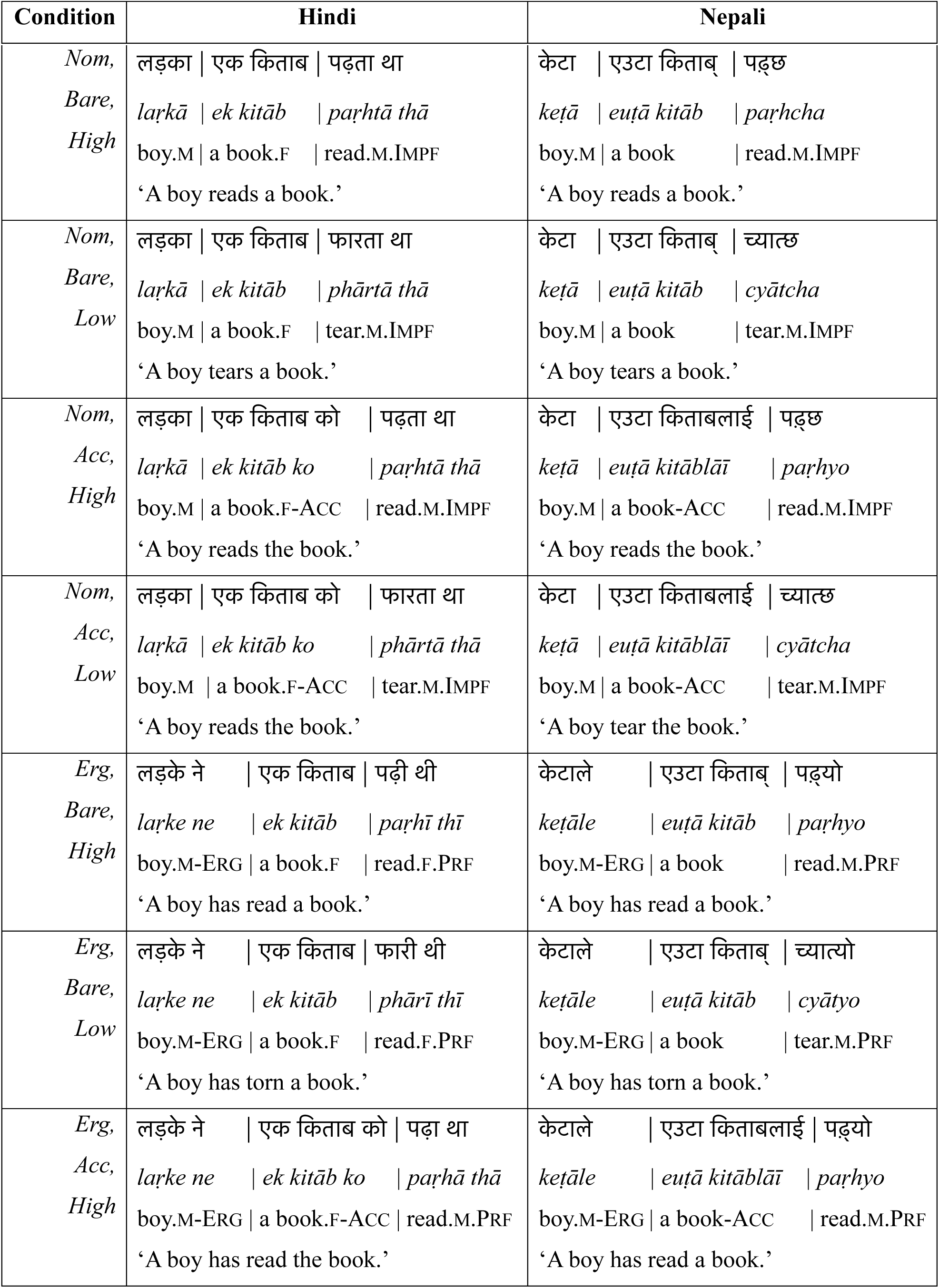
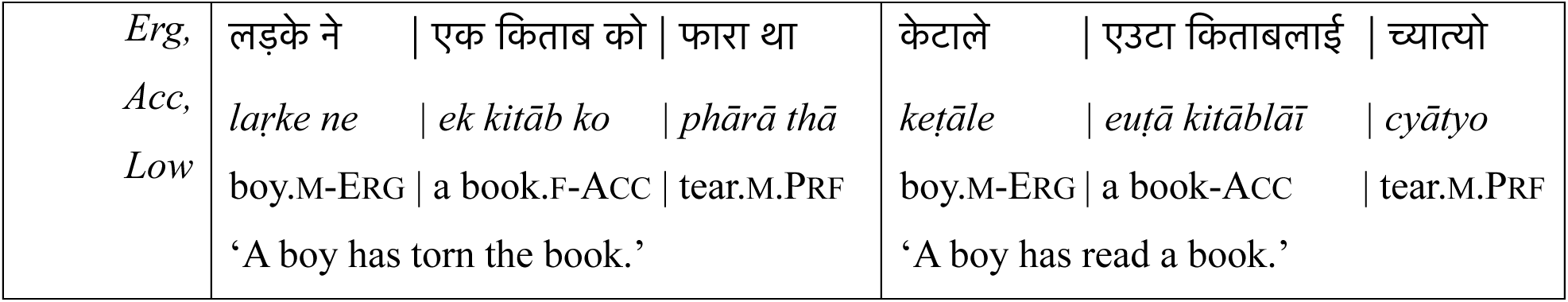
Example stimuli for the experiment, including text in Devanagari script and gloss. Pipes (|) demarcate the three presentation regions: Subject NP, Object NP, and Verb. Gloss abbreviations: ERG = ergative case suffix; ACC = accusative case suffix; M = masculine gender; F = feminine gender; PRF = perfective aspect; IMPF = imperfective aspect.

The subject NPs were animate nouns, and the object NPs were inanimate nouns. This was done to ensure that the intended thematic relations between NPs were easily determinable, to ensure both bare and accusative object NPs were grammatically well-formed and identifiable, and to make object-subject-verb (OSV) analyses less likely. The genders of the NPs were counterbalanced, such that the subjects were grammatically masculine in 50% of the items, and grammatically feminine in the other 50%. In Hindi, inanimate NPs are also marked for gender, so we ensured that the object NP and the subject NP had opposite gender marking. In Nepali, inanimate NPs are not grammatically marked for gender, and default to masculine. Thus, in Nepali, we counterbalanced the gender of the subject NP only.

The verb forms were grammatical in all trials. In Hindi, the verbs were marked with the past perfective endings if the subject was ergative, and with the past imperfective endings if the subject was nominative. In Hindi, tense surfaces as a separate auxiliary form, and aspect is marked with a verb stem suffix. Both tense and aspect markers agree with the subject in gender. In Nepali, the verbs were marked with the present perfective endings if the subject was ergative, and with the present imperfective endings if the subject was nominative. Due to orthographic conventions, the ergative and accusative case suffixes were written after a space in Hindi, and were written connected to the noun stem in Nepali. Because some masculine nouns in Hindi have the same form for the singular and plural, we also included the singular indefinite article एक *ek* ’one’ before the object NP in Hindi, and the singular indefinite article एउटा *euṭā* ’one’ in Nepali to maintain parallelism between the languages.

Verb stems were chosen to ensure that the sentences all contained plausible thematic relations between the arguments and the verb. However, we varied the predictability of the verb stem as a factor in our design. This M350/N400 manipulation was included as a verification of the design, to ensure that both participant groups attended to the stimuli, and that we could replicate an otherwise well-attested neural response. We therefore expect to see a similar response that distinguishes between high-probability and low-probability verb stems around 200–400ms after verb onset in the left temporal lobe for both languages. We treat cloze probability as a binary factor, with levels High Cloze Probability and Low Cloze Probability.

Verb stem probability was estimated from an online sentence completion task (N = 76 for Hindi, N = 48 In Nepali). Subject and object NPs were presented in each case combination, distributed in a Latin Square design for an internet-based task. Participants were instructed to type a completion for the prompt, in either Devanagari or Roman script. Verb stems were identified by two linguists, one a native user of both Hindi and Nepali. The average High Cloze Probability stems were produced 38.1±3.0% of the time for Hindi and 36.7±2.5% of the time for Nepali, and the average Low Cloze Probability stems were produced in 5.2±0.1% and 3.7±0.8% of the completions for Hindi and Nepali respectively.

### 2.2 Procedure

Participants were instructed about the task and signed a consent form electronically. After the participants arrived, we digitized the participant’s head shape using a Polhemus FastScan II (Colchester, VA, USA). This digitized head shape was reduced to a basic surface for use in coregistration with MEG sensor data and MRI structurals. This included 8 fiducial points; 3 points on the forehead and 2 preauricular points approximately 1cm in front of the tragus for coregistration with MEG data, and the two tragus points and one nasion point for coregistration with MRI structurals.

After head shape digitization, participants laid supine in a dimly lit, magnetically shielded room (MSR; Vacuumschmelze, Hanover, Germany). Participants read sentences in Hindi or Nepali presented phrase-by-phrase (subject NP, object NP, verb) on a projected screen. The stimuli were presented in Devanagari script in the Devanagari MT font, printed in 36 point white font against a dark grey background. Each phrase was displayed for 900ms, followed by a 100ms blank screen before the next phrase. This presentation time is longer than in previous studies using Devanagari script (Nevins et al. 2007: 400+200ms; Bickel et al. 2015: 650+100ms), which in turn are longer than typical rapid serial visual presentation (RSVP) paradigms (e.g., 300ms+300ms). We selected this longer presentation time because the phrases displayed on the screen spanned between one and three orthographic words. The interstimulus interval of 900ms+100ms was selected as reasonable after pilot runs with the native-speaking authors.

Each sentence was preceded by a fixation cross that lasted for 600ms. After 25% of trials, participants saw a stock photo image and were instructed to indicate whether the stock photo corresponded to the previous sentence using a button box. The stock photos were retrieved from pexels.com, and were selected by the authors to either clearly depict the activity described in the photo for the ‘match’ trials, or clearly depict a different kind of activity for the ‘mismatch’ trials. The photos were high definition, full-color photos, and were stretched to fill the entirety of the screen. ‘Mismatch’ trials involved either clear depiction of the subject NP referent engaging in a different activity than the one described by the verb phrase, a clear depiction of a different agent referent engaging in the activity described by the verb phrase, or a picture of a different agent engaged in a different activity. For instance, the mismatch trial for sentences corresponding to ‘the boy read/tore a book’ showed an image of a male child eating a lollipop. Thus, participants needed to comprehend both subject NP and the verb phrase in order to succeed on the task. No feedback was provided to participants. Stimuli were presented in a fully within-subjects design; each participant saw each trial once. Presentation order was pseudo-randomized in eight blocks, such that each item set appeared once in one condition per block, and such that the number of conditions were the same per block. Participants were instructed to take a short break between each block, and the experimenter communicated with the participant during each break. Trial structure is exemplified in Figure 2. All experimental procedures and communication with subjects were conducted in Standard Hindi, Nepali, or English, as per the participant’s request. The experiment lasted approximately 90 minutes.

**Figure 2.**
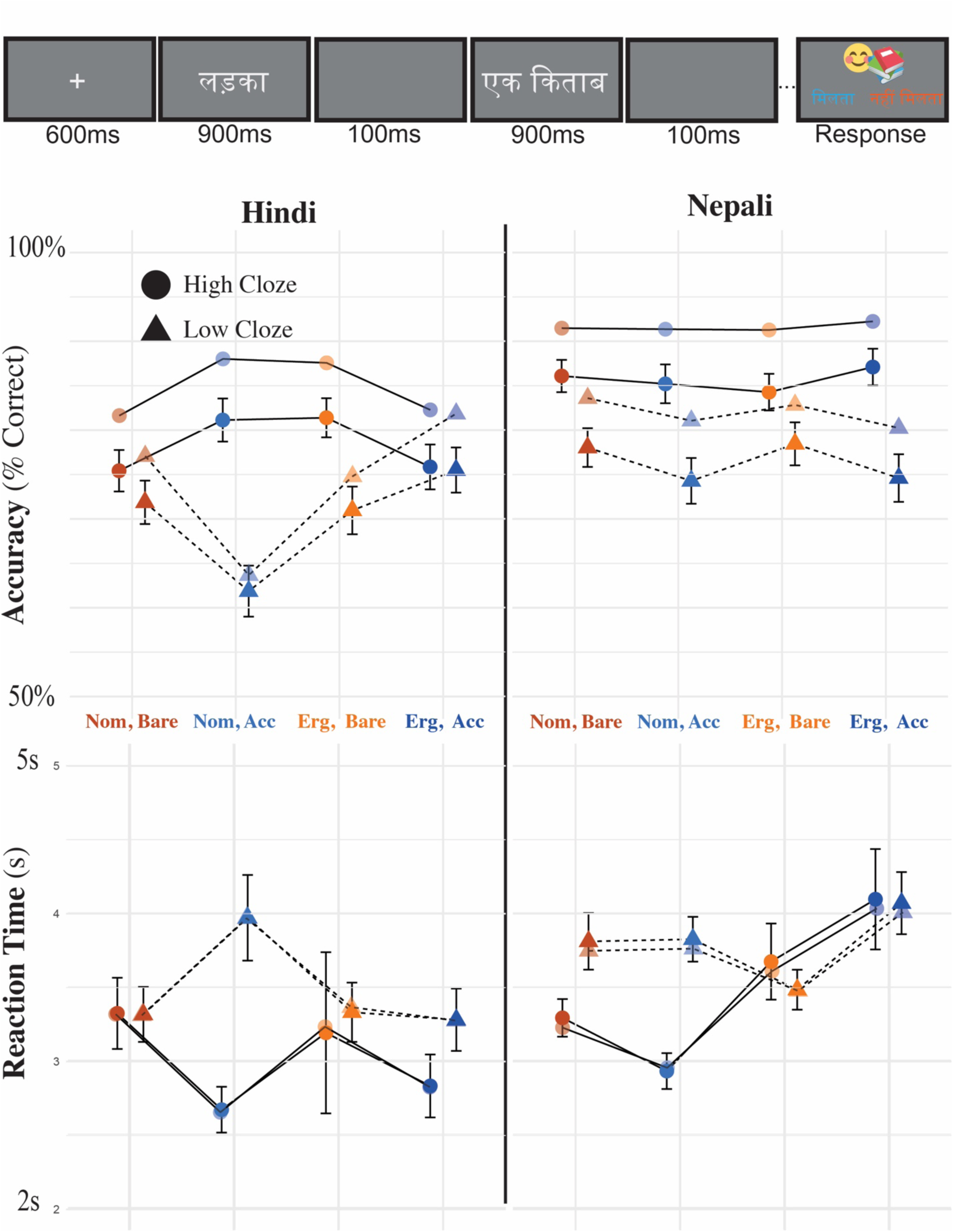
(Top) Task structure. Participants saw a subject NP, object NP, and verb for 900ms each, followed by 100ms of blank screen. Then, they responded whether an image matched the sentence they just read by pressing a key on a keyboard. (Middle and Bottom) Raw and predicted accuracy by condition for accuracty (middle) and reaction time (bottom). Predicted accuracy results are derived from the mixed effects model described in Section 3 and are displayed in a fainter color. Error bars correspond to one standard error above and below the mean.

Brain signals were recorded using a whole-head 208 axial gradiometer (Kanazawa Institute of Technology; Nonoichi, Japan) at a rate of 1000Hz with a 0.1Hz high pass filter and a 200Hz low pass filter applied on-line. The participant’s head position was measured before and after the experiment using the electrical marker coils attached to the two preauricular and three forehead fiducial points.

### 2.3 Data Preprocessing

MEG recordings were noise reduced with the CALM method using the MEG160 software (Adachi et al. 2001). All subsequent data analysis was done in MNE-Python (Gramfort et al. 2013) and Eelbrain (Brodbeck et al. 2023). The data were off-line low-pass filtered at 40Hz using the default settings in MNE-Python. We then used independent component analysis (ICA) to remove periodic neuromagnetic artifacts, including heart beats, saccades, and eye-blinks, and interpolated noisy and flat sensors. We then epoched the sensor data from 200ms prior to the first phrase to the end of the sentence. We rejected all epochs that exceeded a 5 pT peak-to-peak threshold. Baseline correction was applied, using the 200ms pre-stimulus period as baseline.

We then projected the data from sensor space to source space. MEG data were coregistered with either a native T1 MRI structural image (N = 12 for Hindi; *N* = 7 for Nepali) or the FreeSurfer template brain ‘fsaverage’ (CorTechs Labs Inc., California, USA and 175 MGH/HMS/MIT Athinoula A. Martinos Center for Biomedical Imaging, Massachusetts, USA). For participants without a native MRI structural image, the fsaverage template brain was scaled in three dimensions to match the digitized headshape and fiducial markers. An ico-4 source space was then built with 2,562 vertices per hemisphere using the minimum norm estimate algorithm (Hämäläinen & Ilmoniemi 1994). A forward solution was computed using the boundary element model (BEM; Mosher et al. 1999). Channel noise covariance matrices were estimated using the 200ms baseline period before each trial and regularized using the automated method (Engemann & Gramfort 2015). Combining the forward solution and the noise covariance matrices, an inverse solution was computed for each condition for each subject. For computing the inverse solution, we used a free dipole orientation, yielding unsigned values. Finally, the signals were noise normalized in the spatial dimension, yielding a dynamic statistical parameter map (dSPM; Dale et al. 2000). For group-level analyses and identification of regions of interest, participant’s source spaces were warped to a common fsaverage source space, and anatomical regions were referenced using the parcellation atlases provided by fsaverage brain.

### 2.4 Data Analysis

For the MEG data, we conducted three sets of analyses, both for Hindi and Nepali. The first set of analyses sought to identify effects of case morphology in Hindi and Nepali during the processing of the subject and object NP, and the interaction of case/aspect, agreement inflection, and verb stem cloze probability during the processing of the verb period. For these analyses, we conducted whole-brain searches in source space and whole-scalp searches in sensor space (i.e., standardized MEG sensor positions). We conducted these analyses in an ‘N400’ (200–500ms) and ‘P600’ (600–900ms) time window. These analyses were conducted independently in each language.

For the second set of analyses, we defined a variable corresponding to the (expected) gender feature of the verb with levels Masculine and Feminine, given the gender specifications of the subject and object NP, the morphosyntactic case marking on each NP, and the verb agreement rules of each language. In languages like Nepali (or English), the morphological features of the verb are dependent on the morphological features of the subject in grammatical sentences. This means that a brain response to feminine (or plural) verbs might be difficult to distinguish from the brain response to retrieving or activating a feminine (or plural) subject NP, for instance. However, because of the split-ergative agreement system in Hindi, the morphological features of the verb in Hindi are orthogonal to the morphological features of one NP across all sentence types. These analyses were conducted during the processing of the object NP, since this is when comprehenders of both languages had accumulated enough evidence to encode the expected gender feature of the verb. These analyses were also whole-scalp, sensor-space analyses and whole-brain, source-space analyses conducted in the N400 and P600 time windows. The specification of the factor Verb Gender is given in Table 2.

**Table 2.** Assignment of the feature Verb Gender in Hindi and Nepali, as a function of the language, the genders of the subject and object, the subject case, and the object case.

| **Language** | **NP Genders**<br>**(Subject/Object)** | *Subject Case* | *Object Case* | *Verb Gender* |
| --- | --- | --- | --- | --- |
| Hindi | Masculine Subject /<br>Feminine Object | Nominative | Bare | Masculine |
| Hindi | Masculine Subject /<br>Feminine Object | Nominative | Accusative | Masculine |
| Hindi | Masculine Subject /<br>Feminine Object | Ergative | Bare | Feminine |
| Hindi | Masculine Subject /<br>Feminine Object | Ergative | Accusative | Masculine<br>(default) |
| Hindi | Feminine Subject /<br>Masculine Object | Nominative | Bare | Feminine |
| Hindi | Feminine Subject /<br>Masculine Object | Nominative | Accusative | Feminine |
| Hindi | Feminine Subject /<br>Masculine Object | Ergative | Bare | Masculine |
| Hindi | Feminine Subject /<br>Masculine Object | Ergative | Accusative | Masculine<br>(default) |
| Nepali | Masculine Subject | Nominative | Bare | Masculine |
| Nepali | Masculine Subject | Nominative | Accusative | Masculine |
| Nepali | Masculine Subject | Ergative | Bare | Masculine |
| Nepali | Masculine Subject | Ergative | Accusative | Masculine |
| Nepali | Feminine Subject | Nominative | Bare | Feminine |
| Nepali | Feminine Subject | Nominative | Accusative | Feminine |
| Nepali | Feminine Subject | Ergative | Bare | Feminine |
| Nepali | Feminine Subject | Ergative | Accusative | Feminine |

For the third set of analyses, we conducted theoretically-motivated, targeted region-of-interest (ROI) analyses in key source space regions in both languages. This analysis was done to maximize the comparison between the two languages. The ROI analyses sought to identify the same effects as the previous analyses. These analyses were conducted on each language independently, but permit qualitative cross-language comparisons. In this analysis, we identified a cluster showing a significant Subject NP Case:Object NP Case interaction in the left inferior parietal lobe (LIPL) in Hindi. To establish that this effect was specific to Hindi, we then conducted a *post-hoc* analysis in these space and time coordinates across the two languages.

In all analyses, we decided to focus on the N400 and P600 time windows. This was done for a few reasons. First, the epochs in our study are longer than typical in an RSVP task.

Including unnecessary space-time points in a spatio-temporal cluster-based permutation test can increase the false-negative error rate. Secondly, searching the entire epoch runs the risk of finding effects triggered by the differences in morpho-orthographic form which typically occur 200ms and earlier (e.g., Solomyak & Marantz 2011), such as the word length differences between the case-marked and bare NPs. Importantly, we want to separate these processes from the integrative and predictive processes that typically occur later. Moreover, due to the features of the abugida writing system, quantifying variables such as word length for abugidas is not obvious (see Moitra et al. 2024), so regressing out more superficial aspects of the wordforms is not straightforward for this study. Thirdly, initial visual inspection of both source and sensor space data suggested a biphasic response to case in both language groups, centered on both the typical N400 and P600 time window. Lastly, spatio-temporal cluster-based permutation tests do not explicitly test hypotheses about either the latency or the spatial location of an effect. Rather, the clusters identified in spatio-temporal cluster-based permutation tests only establish that there is an effect in the parameters of the search window (Sassenhagen & Draschkow 2019). Thus, whole-epoch and whole-brain searches only establish that there is a significant effect in the epoch or in the brain, whereas search parameters that are constrained *a priori* are more informative. In other words, extending the analysis to the entire time window reduces the certainty in the timing of the effects, which is crucial to the theoretical question.

For the ROI analyses, we averaged the brain response of three critical regions implicated in previous studies on case and agreement morphology – LIFL, LPTL, and LIPL. These ROIs were chosen because activity in these areas have been observed in many previous studies on morphological case processing (including in ergative languages), agreement processing, and syntactic processing, as reviewed in Section 1.1.

These ROIs were defined using the PALS_B12_Brodmann atlas and the aparc_sub atlas provided with the fsaverage template brain. The fsaverage sources corresponding to the left Brodmann Areas (BA) BA-44, BA-45, BA-46, and BA-47 according to PALS_B12_Brodmann parcellation were combined into one label and selected for the LIFL ROI. We used a similar procedure for the sources corresponding to middle temporal 1–4, superior temporal 1–6, transverse temporal 1–2, and banks sts 1–3 to construct the LPTL ROI, and for the sources corresponding to BA-39 and BA-40 in PALS_B12_Brodmann for the LIPL ROI.

For whole brain and ROI analyses, we conducted (spatio)temporal cluster-based permutation tests for null hypothesis testing, following the procedure described by Maris & Oostenveld (2007). For the analyses conducted in the subject NP position, we conducted 1-way ANOVAs contrasting the levels of Subject Case (Ergative, Nominative) at each time point and source (whole brain analyses) and at each time point of the averaged ROI signal (ROI analyses). For analyses conducted in the object NP position, we conducted 2 × 2 ANOVAs with factors Subject Case (Ergative, Nominative), Object Case (Accusative, Bare), and their interaction term, and separately, we conducted 1-way ANOVAs with factor Verb Gender (Masculine, Feminine). Lastly, for the verb position, we conducted 2 × 2 × 2 ANOVAs in the whole-brain analysis, Verb Cloze Probability (High, Low) × Aspect (Perfective, Imperfective) × Gender (Masculine, Feminine). The factor Aspect was a recoding of the factor Subject Case (Ergative=Perfective; Nominative=Imperfective), and the factor Gender was the same as the Verb Gender factor described in Table 2. We included these three factors because brain activity elicited by Low and High Verb Cloze Probabilities verbs may overlap in space or time with brain activity corresponding to verb aspect or gender. However, for the ROI analysis on verb cloze probability during the verb position, we only included the LPTL, since posterior portions of superior and middle temporal gyrus are normally implicated in M350/N400 effects (Lau et al. 2008). For the ROI analyses, we only conducted 1-way ANOVAs of Verb Cloze Probability, to simplify the interpretation of the results.

For the clustering procedure, we identified clusters of adjacent source and time points (whole brain analyses) or time points (ROI analyses) with uncorrected *p*-values < 0.05. We then repeated this procedure 10,000 times per analysis, randomly permuting the labels before calculating the *F*-values and identifying the clusters. This produces a bootstrapped null distribution against which to compare the original clusters; corrected *p*-values correspond to the original cluster’s rank in this null distribution.

For each coefficient of the ANOVA, we used the Benjamin-Hochberg method to correct for false discovery rate (FDR; Benjamini & Hochberg 1995). For FDR correction, we included all (spatio)temporal clusters with *p*-value < 0.10. When we did not identify a significant cluster for a coefficient in the ANOVAs for some time window-ROI search, we included in the FDR correction procedure the a *p*-value of 1.0 if no cluster was identified. For FDR correction, we separate the analyses by language, but correct across time window (N400, P600 time window) and location (hemisphere; ROI). We report on the largest significant cluster in sensor space and targeted ROI analyses, and the largest significant or marginally significant cluster in each hemisphere for the whole brain analyses. We included both hemispheres separately because the adjacency matrix for the fsaverage brain does treat left and right hemisphere as adjacent, and thus brain activity that spans both hemispheres would be reported as two separate clusters.

## 3. Results

### 3.1 Behavioral Results

For behavioral results, we fit a logistic linear mixed effects model in R to the accuracy and a linear mixed effects model to the reaction times on the trials with the picture matching task (R Core Team 2021; Bates et al. 2015). In both analyses, we fit two separate models to the Hindi and Nepali data, because models including a factor of Language did not converge, even including the simplest random effects structure. Each of the four models were fit with fixed effects for Subject Case, Object Case, and Verb Stem Cloze Probability. These factors were sum coded, with 1 for Ergative, Accusative, and Low Cloze Probability respectively, and –1 for Nominative, Bare, High cloze Probability. We fit a ‘maximal’ random effect structure (Barr et al. 2013), including random slopes for Subject Case, Object Case, Verb Stem Cloze Probability and their interactions by participant and by Item. These models did not initially converge, so we used backwards elimination of terms in the random effect structure using likelihood ratio tests until the model converged. In all cases, this procedure resulted in a model with random intercepts for participant and item only. The summary of these findings are presented in Table 3.

**Table 3.** Results of mixed effects models fit to the accuracy and reaction time data for the picture matching task. The structure of each model was: Correct/Reaction Time ∼ Subject Case * Object Case * Verb Cloze Probability + (1|Participant) + (1|Item). Bold mark *p*-values < 0.05.

| **Hindi Acc** | $\beta$ | SE | $z$ | $p$ | **Nepali Acc** | $\beta$ | SE | $z$ | $p$ |
| --- | --- | --- | --- | --- | --- | --- | --- | --- | --- |
| *(Intercept)* | 1.42 | 0.28 | 4.97 | < **0.01** | *(Intercept)* | 1.95 | 0.21 | 9.32 | < **0.01** |
| *SubjC* | –0.10 | 0.14 | –0.75 | 0.45 | *SubjC* | 0.00 | 0.20 | 0.01 | 0.99 |
| *ObjC* | 0.02 | 0.14 | 0.14 | 0.89 | *ObjC* | 0.03 | 0.20 | 0.15 | 0.88 |
| *VCloze* | 0.33 | 0.06 | 5.61 | < **0.01** | *VCloze* | 0.44 | 0.06 | 7.01 | < **0.01** |
| *SC:OC* | 0.02 | 0.14 | 0.14 | 0.89 | *SC:OC* | 0.02 | 0.20 | 0.09 | 0.93 |
| *SC:VC* | 0.10 | 0.06 | 1.78 | 0.08 | *SC:VC* | –0.02 | 0.06 | –0.40 | 0.69 |
| *OC:VC* | –0.04 | 0.06 | –0.69 | 0.49 | *OC:VC* | –0.06 | 0.06 | –0.93 | 0.35 |
| *SC:OC:VC* | –0.25 | 0.06 | –4.30 | < **0.01** | *SC:OC:VC* | 0.02 | 0.06 | 0.28 | 0.77 |
| **Hindi RT** | $\beta$ | SE | $t$ | $p$ | **Nepali RT** | $\beta$ | SE | $t$ | $p$ |
| *(Intercept)* | 3.25 | 0.37 | 8.78 | < **0.01** | *(Intercept)* | 3.60 | 0.29 | 12.58 | < **0.01** |
| *SubjC* | 0.07 | 0.14 | 0.51 | 0.61 | *SubjC* | –0.18 | 0.15 | –1.18 | 0.24 |
| *ObjC* | 0.06 | 0.14 | 0.46 | 0.64 | *ObjC* | –0.09 | 0.15 | –0.57 | 0.57 |
| *VCloze* | –0.24 | 0.10 | –2.48 | **0.01** | *VCloze* | –0.15 | 0.06 | –2.23 | **0.03** |
| *SC:OC* | –0.06 | 0.14 | –0.45 | 0.65 | *SC:OC* | 0.05 | 0.07 | 0.75 | 0.45 |
| *SC:VC* | –0.09 | 0.10 | –0.95 | 0.34 | *SC:VC* | –0.19 | 0.07 | –2.84 | < **0.01** |
| *OC:VC* | 0.20 | 0.10 | 2.15 | **0.03** | *OC:VC* | 0.05 | 0.07 | 0.75 | 0.45 |
| *SC:OC:VC* | 0.13 | 0.10 | 1.32 | 0.19 | *SC:OC:VC* | 0.02 | 0.07 | 0.34 | 0.73 |

For both Hindi and Nepali participants, responses were more accurate and faster to sentences with high probability verbs compared to low probability verbs. We also observed a three-way interaction effect of Subject Case, Object Case, and Verb Stem Cloze Probability in Hindi participants, showing less accurate responses for sentences with ergative subjects, accusative objects, and low probability verbs. This was not observed in the Nepali participants. However, we observed a two-way interaction effect of Subject Case and Verb Stem Cloze Probability in Nepali-speaking participants, showing slower responses for nominative subjects with low probability verbs. The mean acceptability and response times, and the estimates of these linear models, are shown in Figure 2.

### 3.2 Whole-Brain Source Space Results

We first discuss the whole-brain source space analyses, then the whole-scalp sensor-space analyses. Because the effects of case morphology and the encoding of agreement features during the object NP period are the most directly relevant to our hypothesis, we discuss these before discussing the effects observed during the subject NP and verb periods.

#### Object NP Period

In the whole-brain analyses in Hindi, we observe a Subject NP Case cluster in the N400 time window (Fig 3B, 200–500ms, adjusted *p* = 0.08), centered primarily on LATL, and Subject NP Case cluster in the P600 time window (Fig 3C, 600–899ms, adjusted *p* = 0.20). This cluster is centered on RPTL and RIPL. Both clusters are not significant after correction for multiple comparisons, although the first cluster was significant before correction. In *post-hoc* pairwise comparisons, the earlier cluster shows greater activity for bare object NPs after an ergative subject NP.

**Figure 3.**
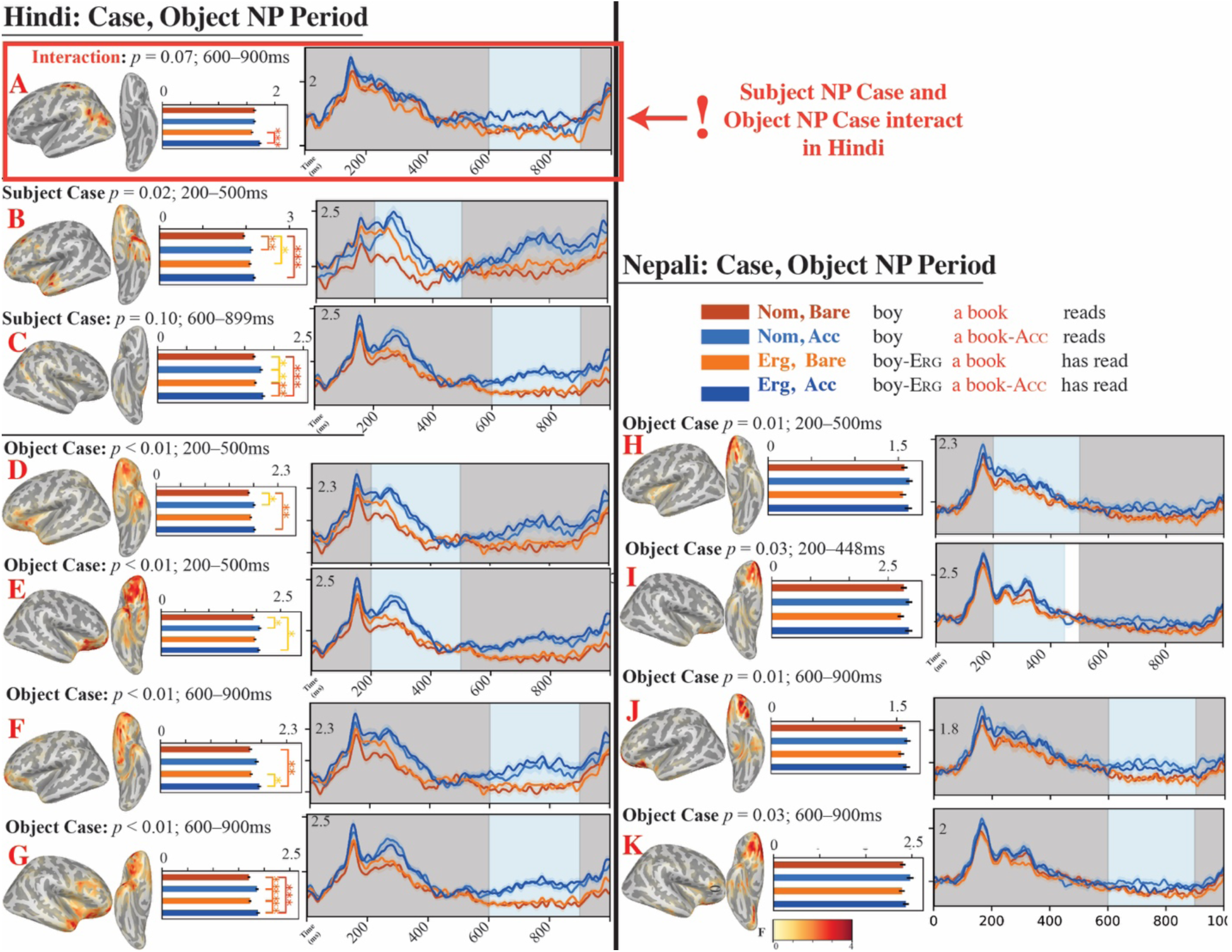
Results of whole-brain spatio-temporal cluster-based permutation tests for Subject NP Case × Object NP Case ANOVAs during the object NP period in Hindi and Nepali. Brain plots show the spatial extent of the spatio-temporal cluster and the *F*-values, averaged over time. Bar plots show activation (dSPM) in the cluster, averaged over space and time; starred comparisons show significant (* *p <* 0.05; ** *p* < 0.01) pairwise comparisons using *t*-tests with Hochberg correction. Time courses show averaged activity (dSPM) over time (ms) in the spatial coordinates of the cluster. Gray shading indicates time points that were excluded from the search space, and blue shading indicates the temporal extent of the cluster. Each cluster is reported with its coefficient, uncorrected *p*-value, and temporal extent.

In Hindi, four Object Case clusters are observed during the object NP period. During the N400 time window, two clusters were centered on the LATL, RATL, and OF (Fig 2D, 200– 500ms, adjusted *p* = 0.04; Fig 3E, 200–500ms, adjusted *p* = 0.04), potentially indexing bilateral brain activity. Both clusters show greater activation for the accusative-marked object NPs compared to the bare object NPs. A similar pattern was observed in the two clusters identified in the P600 time window (Fig 3F, 600–900ms, adjusted *p* = 0.02; Fig 3G, 600–900ms, adjusted *p* = 0.02), centered over bilateral ATL, OF, and right inferior frontal lobe (RIFL).

Finally, we observed one cluster for the Subject Case × Object Case interaction during the P600 time window (Fig 3A, 600–900ms, adjusted *p* = 0.28), centered over LPTL, occipito-temporal lobe, and LIFL. This cluster was marginally significant before correction. This cluster showed greater activity for Ergative Subject NPs, Accusative Object NPs trials (NP*-ne* NP–*ko*; default agreements structures) over Ergative Subject NPs, Bare Object NPs (NP-*ne* NP; object agreement structures), but did not show a similar pattern for Nominative Subjects (NP NP, NP NP-*ko*; both subject agreement structures). This cluster demonstrated the interaction between morphosyntactic case features that we predicted for Hindi.

In Nepali, we did not observe any Subject NP Case clusters that met our criteria. Two Object NP Case clusters were observed in the N400 time window, centered on the left and right OF (Fig 2H, 200–500ms, adjusted *p*-value = 0.04; Fig 2I, 200–448ms, adjusted *p*-value = 0.04). As in Hindi, these clusters show greater activity for accusative-marked object NPs, and likely depict a single bilateral OF response. Also similar to Hindi, there were two Object NP Case clusters in the P600 time window (Fig 3J, 600–900ms, adjusted *p*-value = 0.04; Fig 3K, 600– 900ms, adjusted *p*-value = 0.04), centered on left and right OF. As in Hindi, this likely demonstrates a sustained increase in bilateral OF activity for accusative-marked object NPs. There were no Subject NP Case × Object NP Case interaction clusters.

For the predicted verb gender analysis, in the whole-brain analyses in Hindi, we observed two clusters in the N400 time window and one cluster in the P600 time window during the object NP period. During the N400 time window, both Verb Gender clusters in the left hemisphere (Fig 3D, 200–500ms, adjusted *p*-value = 0.04) and the right hemisphere (Fig 3E, 200–500ms, adjusted *p*-value = 0.04) showed greater activity for (expected) masculine gender verbs compared to (expected) feminine gender verbs. Both clusters were centered on OF and the insula, and the left hemisphere cluster also included portions of the LATL and LIFL. In the P600 time window, we identified a Verb Gender cluster (Fig 3F, 600–900ms, adjusted *p*-value = 0.08), significant prior to correction, that showed a continued higher activation for expected masculine gender verbs in the left OF. Taken together, these results likely identify a persistent bilateral response in anterior frontal and temporal lobe during the object NP time window distinguishing between upcoming masculine and feminine verbs. This is consistent with our hypothesis that Hindi comprehenders encode an expectation for the morphosyntactic features of the verb after initial processing of the object NP. In Nepali, we did not identify any Verb Gender clusters that were consistent with our criteria.

#### Subject NP Period

In Hindi, during the subject NP time period, one Subject Case cluster was observed in the N400 time window (Fig 4A, 200–500ms, adjusted *p*-value = 0.11), centered over the ventral surface of temporal and frontal lobe. This cluster was marginally significant prior to correction. It shows greater activity for ergative-marked subject NPs compared to nominative-marked subject NPs. In the P600 time window, we observed two significant Subject NP clusters (Fig 4B, 600–900ms, adjusted *p* = 0.04; Fig 4C, 600–900ms, adjusted *p* = 0.04). These clusters are centered on ventral surface of the frontal lobe in both left and right hemispheres, and show greater activity for ergative-marked subject NPs compared to nominative-marked subject NPs. This likely shows a single bilateral increase in activity in this region.

**Figure 4.**
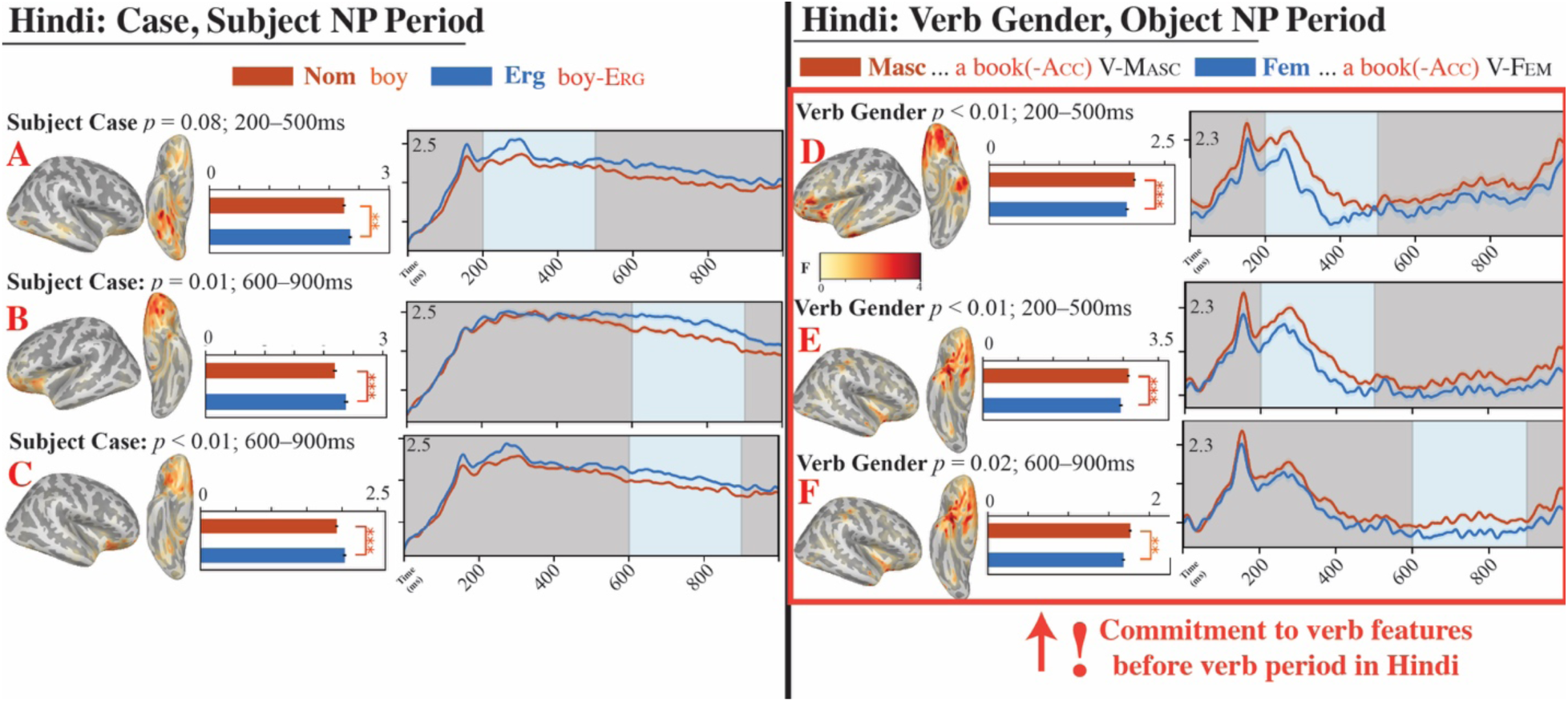
Results of whole-brain spatio-temporal cluster-based permutation tests for Subject NP and Verb Gender ANOVAs during the object NP period in Hindi. Brain plots show the spatial extent of the spatio-temporal cluster and the *F*-values, averaged over time. Bar plots show activation (dSPM) in the cluster, averaged over space and time; starred comparisons show significant (* *p <* 0.05; ** *p* < 0.01) pairwise comparisons using *t*-tests with Hochberg correction. Time courses show averaged activity (dSPM) over time (ms) in the spatial coordinates of the cluster. Gray shading indicates time points that were excluded from the search space, and blue shading indicates the temporal extent of the cluster. Each cluster is reported with its coefficient, uncorrected *p*-value, and temporal extent. No clusters were observed in Nepali.

No Subject NP clusters were identified in Nepali that met our criteria.

#### Verb Period

The goal of analyzing the verb period is to have an independent verification that comprehenders interpret the sentence. We expect a greater M350 effect for low cloze probability verb stems compared to other verb stems. However, because we could not anticipate that the effects of the factor Verb Stem Cloze Probability would necessarily be independent in space and time from the morphological Aspect or Verb Gender, we included these factors as covariates. We only report on main effects of Verb Stem Cloze Probability and its interaction terms, since these are most relevant for the independent verification. The neural responses to gender marking and aspect during the processing of the verb are not necessarily critical to the hypotheses that we test here, and therefore we do not interpret them here. We first report main effects of Verb Stem Cloze Probability, followed by interactions with Verb Gender, then interactions with Aspect.

In Hindi, we did not identify any clusters of Verb Stem Cloze probability during the N400 time window. During the P600 time window, we found a cluster in the left insula and LIFL (Fig 5A, 600–900ms, adjusted *p* = 0.16), significant prior to correction. Verb Stem Cloze Probability × Verb Gender interaction clusters were observed in right superior parietal regions (Fig 5B, 600– 900ms, adjusted *p* = 0.06) and left superior temporal lobe and LMTL during the P600 time window (Fig 5C, 618–900ms, adjusted *p* = 0.10). Both clusters demonstrate a modulation of the Verb Stem Cloze probability effect by the gender specification on the verb. Finally, Verb Stem Cloze Probability × Verb Aspect clusters were observed in RIPL (Fig 5D, 200–500ms, adjusted *p* = 0.08) and LMFL (Fig 5E, 200–500ms, adjusted *p* = 0.08) during the N400 time window, and right occipito-parietal and OF regions during the P600 time window (Fig 5F, 614–900ms, adjusted *p* = 0.08). All reported clusters in Hindi were marginally significant either before or after correction for multiple comparisons.

**Figure 5.**
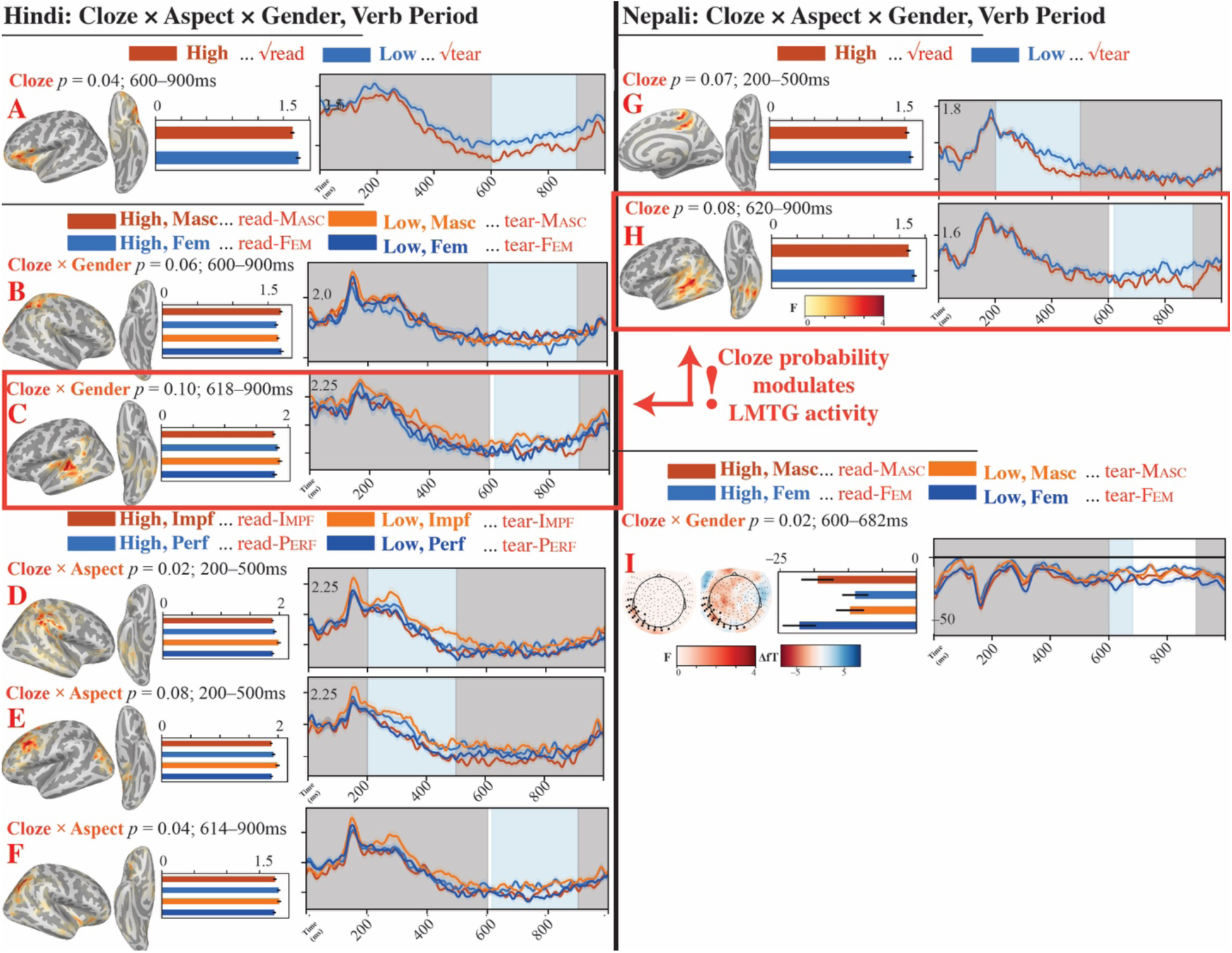
Results of whole-brain and whole-scalp spatio-temporal cluster-based permutation tests for Verb Stem Cloze Probability × Aspect × Verb Gender ANOVAs during the verb period in Hindi and Nepali. Brain plots show the spatial extent of the spatio-temporal cluster and the *F*-values, averaged over time. Topographic plots show the spatial extent of the spatio-temporal cluster and the *F*-values averaged over time (left) and the topographic plot of the difference (fT) between Low and High Verb Stem Cloe Probability (right). Bar plots show activation (dSPM; fT) in the cluster, averaged over space and time; starred comparisons show significant (* *p <* 0.05; ** *p* < 0.01) pairwise comparisons using *t*-tests with Hochberg correction. Time courses show averaged activity (dSPM) over time (ms) in the spatial coordinates of the cluster. Gray shading indicates time points that were excluded from the search space, and blue shading indicates the temporal extent of the cluster. Each cluster is reported with its coefficient, uncorrected *p*-value, and temporal extent.

In Nepali, we identified a cluster in the right premotor cortex and medial parietal surfaces during the N400 time window (Fig 5G, 200–500ms, adjusted *p* = 0.28), and a cluster in LMTL during the P600 time window (Fig 5H, 620–900ms, adjusted *p* = 0.28). Both clusters demonstrated greater activity for Low vs. High Verb Stem Probability verbs, and were marginally significant prior to correction.

### 3.2 Whole-Head Sensor Space Results

#### Object NP Period

For the case morphology analyses during the object NP period, we observed two Object NP Case clusters in the N400 and P600 time window in both Hindi and Nepali. In Hindi, the N400 Object NP Case cluster (Fig 6A, 209–482ms, adjusted *p*-value = 0.02) showed greater positive activity for accusative-marked object NPs compared to bare object NPs in anterior and central sensors, and right laterior and posterior sensors. The P600 Object NP Case cluster (Fig 6B, 600–900ms, adjusted *p*-value = 0.02) showed greater negative activity for accusative-marked object NPs compared to bare object NPs in anterior and central sensors, and left posterior sensors. In Nepali, the N400 Object NP Case cluster (Fig 6C, 207–500ms, *p*-value = 0.02) showed greater positive activity for accusative-marked object NPs compared to bare object NPs in anterior, central, and left posterior sensors. The P600 Object NP Case cluster (Fig 6D, 663– 900ms, adjusted *p*-value = 0.02) showed greater negative activity for accusative-marked object NPs compared to bare object NPs in anterior and central sensors. No clusters were identified according to our criteria for Subject Case in Hindi or Nepali during the object NP period, nor for the Subject Case × Object Case interaction.

**Figure 6.**
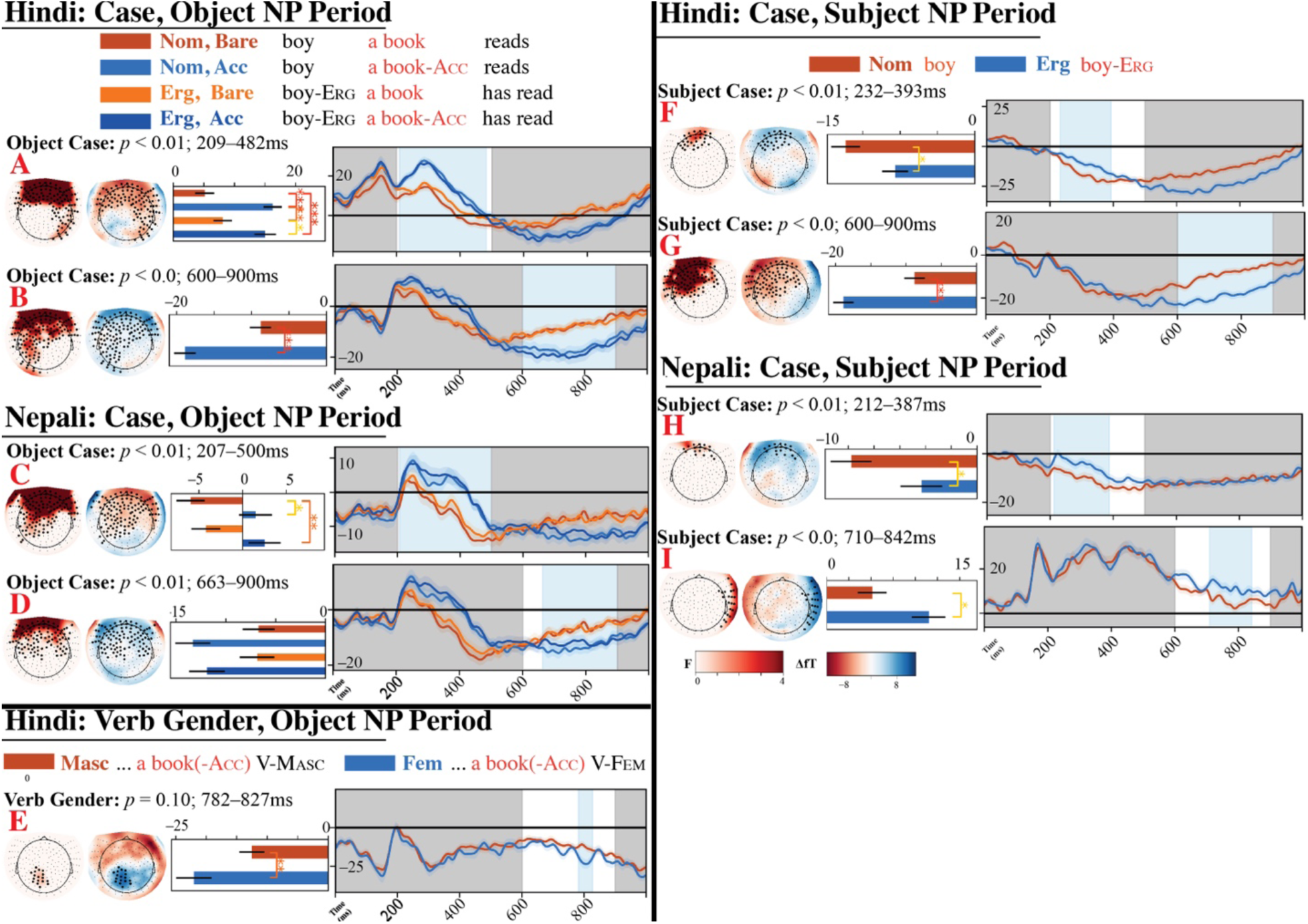
Results of whole-scalp spatio-temporal cluster-based permutation tests for Subject NP, Subject NP Case × Object NP Case, and Verb Gender ANOVAs during the subject and object NP period in Hindi and Nepali. Topographic plots show the spatial extent of the spatio-temporal cluster and the *F*-values averaged over time (left) and the topographic plot of the difference (fT) between the ‘marked’ conditions (Ergative, Acsusative, Feminine) and the ‘unmarked’ conditions (Nominative, Bare, Masculine; right). Bar plots show activation (dSPM) in the cluster, averaged over space and time; starred comparisons show significant (* *p <* 0.05; ** *p* < 0.01) pairwise comparisons using *t*-tests with Hochberg correction. Time courses show averaged activity (dSPM) over time (ms) in the spatial coordinates of the cluster. Gray shading indicates time points that were excluded from the search space, and blue shading indicates the temporal extent of the cluster. Each cluster is reported with its coefficient, uncorrected *p*-value, and temporal extent.

In the whole-scalp sensor-space analyses, we also identified one cluster in the P600 time window in Hindi for Verb Gender (Fig 6E, 782–827ms, adjusted *p*-value = 0.20), that was marginally significant prior to correction. This cluster showed a short period of greater negative activity for expected feminine verbs compared to masculine verbs, centered over central and posterior sensors. In Nepali, we did not identify any Verb Gender clusters that were consistent with our criteria.

#### Subject NP Period

In the whole-head sensor-space analyses, we observe Subject NP Case clusters during the N400 and P600 time window in subject NP period in both Hindi and Nepali. In Hindi, the earlier cluster (Fig 6F, 232–393ms, adjusted *p-*value = 0.02) shows greater negative activity for nominative-marked subject NPs over ergative-marked subject NPs in anterior midline sensors.

The later P600 cluster (Fig 6G, 600–900ms, adjusted *p*-value = 0.02) shows the opposite pattern, with greater negative activity for ergative-marked subject NPs over nominative-marked subject NPs in anterior, left lateral, and left midline sensors. In Nepali, the N400 cluster (Fig 6H, 212– 387ms, adjusted *p*-value = 0.02) showed greater negative activity for nominative-marked subject NPs compared to ergative-marked subject NPs over anterior sensors, much like in Hindi. The later P600 cluster (Fig 6I, 710–842ms, adjusted *p*-value = 0.02) showed greater positive activity for ergative-marked subject NPs compared to nominative-marked subject NPs over right lateral sensors. Although the two Subject Case clusters observed during the subject NP time window differed in terms of the sensors in which they were statistically significant and waveform morphology, inspection of the topographic plots may suggest that these are similar responses, with greater positive activity observable in the left lateral sensors and greater negative activity in the right lateral sensors.

#### Verb Period

Finally, during the verb period, we observed a Verb Stem Cloze Probability × Verb Gender cluster in Nepali during the P600 time window (Figure 5I, 600–682ms, adjusted *p*-value = 0.08), significant prior to correction. No other clusters were identified according to our criteria in Hindi, nor during the N400 time window in Nepali.

### 3.3 Targeted ROI Analyses

No clusters were observed during the subject NP period in the three ROIs.

For the object NP period, we observed temporal clusters of Object Case in Hindi in the N400 time window in LIPL (Fig 7A, 460–498ms, adjusted *p*-value = 0.21), LPTL (Fig 7B, 256– 326ms, adjusted *p*-value = 0.12), and a marginally significant cluster, pre-correction, in the P600 time window in LIFL (Fig 7C, 706–722ms, adjusted *p*-value = 0.21). The LPTL cluster showed greater activity for the bare object NPs, whereas the LIFL clusters both showed greater activity for the accusative-marked object NPs. We also identified a marginally significant pre-correction Subject Case × Object Case cluster in Hindi in the LIPL (Fig 7D, 768–498ms, adjusted *p*-value = 0.12). This cluster showed a reduction in activity for Ergative Subject NP, Bare Object NP conditions in Hindi. No Subject Case clusters were identified in Hindi that met our criteria, and no clusters were identified in Nepali that met our criteria.

**Figure 7.**
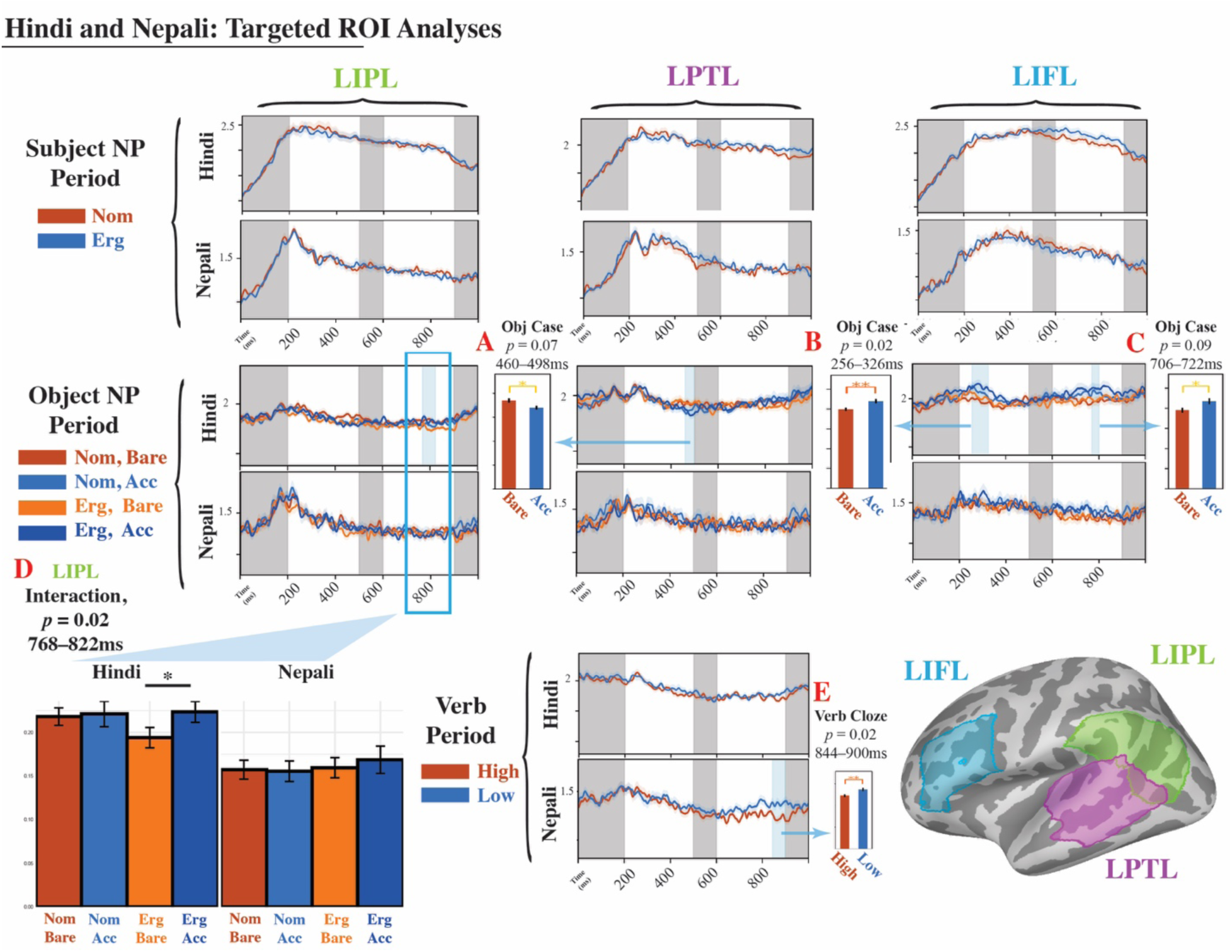
Results of targeted ROI analysis. Columns show averaged activity (dSPM) over time (ms) in the left inferior parietal lobe (LIPL), the left posterior temporal lobe (LPTL), and the left inferior frontal lobe (LIFL) ROIs. Columns correspond to the analyses conducted in the subject NP period (Subject NP Case), object NP period (Subject NP Case × Object NP Case), and verb (Verb Stem Cloze Probability). Gray shading indicates time points that were excluded from the temporal cluster-based permutation tests. Light blue shading indicates the temporal extent of reported clusters. Bar graphs show the activity in the cluster averaged over time, stars indicate significant pairwise comparisons using *t*-tests with Hochberg correction. (Bottom left inset) Comaprison of averaged activity from 768–822ms in Hindi and Nepali in the LIPL, from the Subject NP Case × Object NP Case ROI analysis conducted in the LIPL in Hindi. (Bottom right inset) LIFL, LIPL, and LPTL masks plotted on the fsaverage template brain.

In the Verb Stem Cloze Probability analysis, we identified a cluster in Nepali during the P600 time window in the LPTL ROI (Fig 7E, 844ms–900ms, adjusted *p*-value = 0.02).

To further test the claim that the interactions we’ve observed in Hindi are absent in Nepali, we conducted a *post-hoc* analysis in the LIPL ROI cluster across both Hindi and Nepali participants. For each participant and condition, we averaged the activity (dSPM) in the LIPL during the temporal cluster identified in the ROI analysis (768–822ms). Afterwards, we constructed a linear mixed-effects model using lme4 in R on the average values, with Subject Case, Object Case, and Language as their fixed effects, plus their interaction terms. These factors were sum coded, with 1 for Ergative, Accusative, and Hindi respectively, and –1 for Nominative, Bare, and Nepali. We also included random slopes for these terms by Subject. This model did not initially converge, so we eliminated terms in the random effects structure using backwards elimination with likelihood ratio tests until the model converged. This resulted in a random effect structure with random intercept for participant only. Afterwards, we conducted a pairwise comparison between the levels of Object Case nested within each combination of factors Subject Case and Language, to test the hypothesis that brain activity during the processing of bare and accusative object NPs differed in a way unique to Hindi after an ergative subject NP, in contrast to bare and accusative object NPs after ergative subject NPs in Nepali, or nominative subject NPs in Hindi or Nepali. The final model, the model results, and the pairwise comparison results are given in Table 4.

**Table 4.**
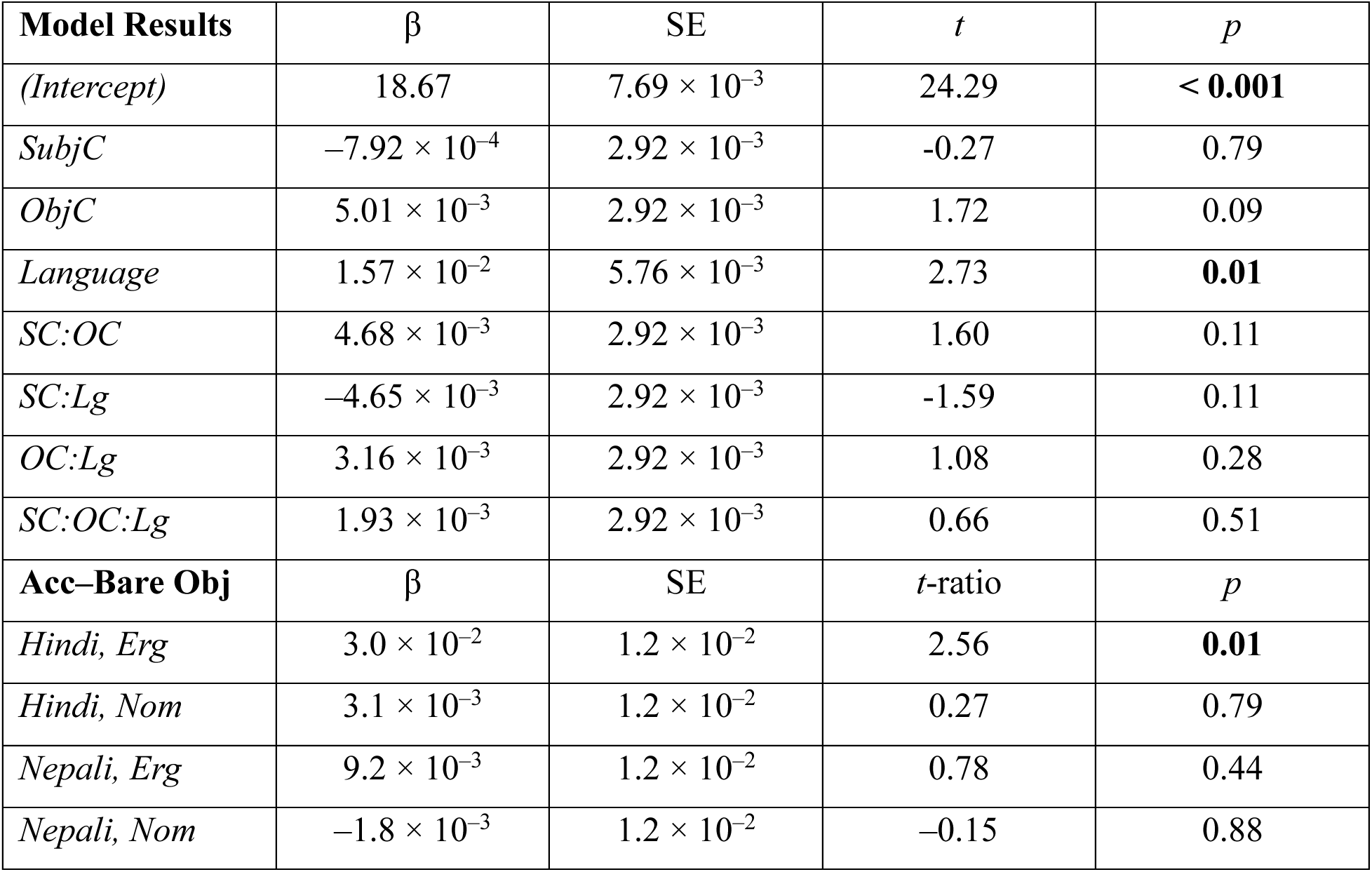
Results of mixed effects models fit to the average activation (dSPM) in the left temporo-parietal junction ROI (Brodmann Areas 39+40), 768–822ms. The structure of each model was: Correct/Reaction Time ∼ Subject Case * Object Case * Verb Cloze Probability + (1|Participant) + (1|Item). Bold mark *p*-values < 0.05.

| **Model Results** | $\beta$ | SE | $t$ | $p$ |
| --- | --- | --- | --- | --- |
| *(Intercept)* | 18.67 | $7.69 \times 10^{-3}$ | 24.29 | **< 0.001** |
| *SubjC* | $-7.92 \times 10^{-4}$ | $2.92 \times 10^{-3}$ | -0.27 | 0.79 |
| *ObjC* | $5.01 \times 10^{-3}$ | $2.92 \times 10^{-3}$ | 1.72 | 0.09 |
| *Language* | $1.57 \times 10^{-2}$ | $5.76 \times 10^{-3}$ | 2.73 | **0.01** |
| *SC:OC* | $4.68 \times 10^{-3}$ | $2.92 \times 10^{-3}$ | 1.60 | 0.11 |
| *SC:Lg* | $-4.65 \times 10^{-3}$ | $2.92 \times 10^{-3}$ | -1.59 | 0.11 |
| *OC:Lg* | $3.16 \times 10^{-3}$ | $2.92 \times 10^{-3}$ | 1.08 | 0.28 |
| *SC:OC:Lg* | $1.93 \times 10^{-3}$ | $2.92 \times 10^{-3}$ | 0.66 | 0.51 |
| **Acc–Bare Obj** | $\beta$ | SE | $t$-ratio | $p$ |
| *Hindi, Erg* | $3.0 \times 10^{-2}$ | $1.2 \times 10^{-2}$ | 2.56 | **0.01** |
| *Hindi, Nom* | $3.1 \times 10^{-3}$ | $1.2 \times 10^{-2}$ | 0.27 | 0.79 |
| *Nepali, Erg* | $9.2 \times 10^{-3}$ | $1.2 \times 10^{-2}$ | 0.78 | 0.44 |
| *Nepali, Nom* | $-1.8 \times 10^{-3}$ | $1.2 \times 10^{-2}$ | -0.15 | 0.88 |

In the full linear model, the only coefficient that was significant was the Language, with significantly more activity in the LIPL in Hindi compared to Nepali. In the pairwise comparisons, we found that there was greater activity for accusative object NPs compared to bare object NPs in Hindi after an ergative subject but not in Nepali, and that there were no significant differences in activity for accusative object NPs compared to bare object NPs after either subject NP type in Nepali.

## 4. Discussion

In both Hindi and Nepali, we found a consistent profile showing correlations between activity in LIFL, LATL, and OF and morphosyntactic case. Additionally, during the processing of the object NP in Hindi, there was a correlation between activity in these regions and the gender specification of the upcoming verb beginning at 200ms, as required by the Hindi split-ergative agreement system. This is consistent with the hypothesis that Hindi comprehenders use the inflectional features of the subject and object NPs as a cue to pre-encode the inflectional features of the verb. No such finding was observed in Nepali. Moreover, we observed that activity in the LIPL showed interactions between the subject and object NP case specifications, both in the whole-brain and targeted ROI analyses. In both analyses, these effects were observed later, emerging at 600ms, and demonstrated distinct patterns. The whole brain analysis revealed a reduction in activity for the Ergative Subject NP, Bare Object NP conditions (NP-*ne* NP; object agreement configuration), and the ROI analysis revealed a reduction in activity for the Ergative Subject NP, Accusative Object NP conditions (NP-*ne* NP*-ko*; default agreement configuration). This activity in the LIPL was also not observed in Nepali.

### Delayed Sensitivity to Cloze Probability

The Verb Cloze Probability manipulation was included as an independent verification that participants were attending to the task and processing the sentences in a similar way. We predicted that the neural response would emerge as a typical M350 response and localize to posterior portions of LMTL.

This prediction was not borne out. During the M350/N400 time window, we observed a typical M350 response in Nepali, but this localized to the medial wall of the right hemisphere. In Hindi, we did not observe a typical M350 response either. Instead, we observed two clusters showing an interaction between aspect and cloze probability in RIPL and LMFL, with greater activity for the less probable imperfective verb. Finally, M350-like effects emerged during the later, P600 time window. In Nepali, the effect emerged in the LMTL, and in Hindi, in the LIFL+insula. Finally, in Hindi, we observed an interaction between Verb Stem Cloze Probability and Gender in LpSTL/LMTL.

Although M350 responses typically localize to LMTL, typical ‘N400’ responses to unpredictable stimuli may also show activity in the LIFL (Lau et al. 2008). Thus, the response to unpredicted verbs in the frontal lobe in Hindi may be consistent with a typical M350/N400 interpretation. However, the delayed response in both languages was surprising. This delay may have been due to the increased complexity of the stimuli compared to stimuli from other N400 studies. In our experiment, the epochs consisted of multiple phonological/orthographic words that were displayed in parallel, which may have led to a delay in the identifiability of a response corresponding to lexical access of the verb stem. Alternatively, the delayed response may reflect additional complexities introduced by the Devanagari writing system. Another potential explanation is that this delay may have reflected an adaptation to the slower-than-usual presentation speed of 1000ms per phrase. Participants may have adjusted their reading strategy, or may have withheld attention at the initial presentation of the word and phrase, thereby delaying stages of lexical access compared to the onset of the stimulus. Finally, ‘N400’ responses to unpredictable stimuli may not be neurobiologically distinct from other responses to unpredictable stimuli that otherwise differ in latency or space (LAN, mismatch negativity (MMN); Bornkessel-Schlesewsky & Schlesewsky 2019); thus, an increase in latency compared to the typical time window may not ‘rule out’ considering these responses as a member of the N400 family, once these other factors are accounted for.

We also note that the cloze probability of our stimuli was overall lower than is typical in N400 studies that categorically contrast brain activity with cloze probability. High cloze probability stimuli usually have completion rates ∼40-50%, and low cloze probability stimuli are closer to ∼5–10%, whereas ours were lower than these. This could have also tempered the robustness of an M350 response that may have otherwise been observed in the N400 time window.

Finally, we note that in both languages, the factor Verb Stem Cloze probability was distributed across different aspect markers and gender agreement. For instance, in Hindi, the Verb Stem Cloze Probability factor collapses across both genders and aspects (High: पढ़ा था *paṛhā thā* read.PERF.MASC, पढ़ी थी *paṛhī thī* read.PERF.FEM, पढ़ता था *paṛhtā thā*, read.IMPF.MASC पढ़ती थी read.IMPF.FEM; Low: फारा था *phārā thā* tear.PERF.MASC, फारी थी *phārī thī* tear.PERF.FEM, फारता था *phārtā thā* tear.IMPF.MASC, फारती थी *phārtī thī* tear.IMPF.FEM). We designed the experiment with the assumption that the predictability of the verb stem would be isolatable as an independent factor, but this may have been erroneous (AUTHORS, in prep.); it may be in principle that certain verb stems may be more or less likely in specific aspects or with certain gender inflections, or that different combinations of NP case markings may alter the predictability of verb stems. Similarly, if similar neuroanatomical regions are engaged in processing the verb stem as well as its morphosyntactic features, then there may be no distinct, separable effects of the verb stem identifiable in the brain.

Other studies in Hindi have demonstrated negative-going ERPs 200–500ms in studies contrasting predictable stimuli vs. violations. Choudhary et al. (2009) report on differences in N400 amplitudes for unlikely subject NP case-verb aspect combinations (ergative-imperfective; nominative-perfective). Bhattamishra et al. (2021) report on N400(-like) responses for verb subject-agreement mismatches between inanimate subject NPs and verbs. Thus, other studies have demonstrated N400 effects for formal inflectional features on the verb given the prior arguments. By contrast, our study focused on the prediction of the morphosyntactic features of the verb, but we did not deploy a prediction violation paradigm. How do previous N400 findings in Hindi relate to “traditional” N400 effects, corresponding to more vs. less predictable verb stems? To our knowledge, no previous study has contrasted neural responses the (un)predictability of the verb stem vs. verb inflection directly, and thus we leave the question of the commensurability of the M350/N400 corresponding to identification of a verb stem and processing of its inflectional features as an open question.

#### Case and Gender in Language Areas

In both Hindi and Nepali, we observed greater activation in the bilateral OF, LATL, LIFL, and RIFL for ergative subjects and accusative objects. These findings could in principle reflect computations corresponding to representing ergative and accusative NPs, respectively. However, both cases are ‘marked’, in the sense that there is an additional, overt morpheme. For this reason, the increased activity in these regions could reflect additional computations corresponding to the visual stimuli, instead of any abstract linguistic computation. Alternatively, these responses may reflect the morphosyntactic processing of the noun into its stem and case ending. The ergative case and accusative case markers also provide additional semantic information about the interpretation of the sentence that may be relevant for predictive processing. In both languages, the ergative case implies that the sentence will terminate with a perfective verb (Choudhary et al. 2009). Similarly, in both languages, the accusative case implies a definite or specific interpretation to the object NP. Thus, activity associated with main effects of case morphology could in principle reflect increased visual processing, morphosyntactic decomposition of the argument noun, top-down application of verb aspect prediction, or updates to the discourse representation, or some combination of these computations. For this reason, we hesitate to draw strong conclusions about these responses, and report them for the sake of completeness and future comparisons with other studies.

We note that the LIFL, LATL, and (L)OF are frequently implicated in fMRI and MEG studies on the processing of complex structures. LIFL has been implicated in many processes in language comprehension. LIFL activity is often highly correlated with general processes assisting in sentence comprehension (Stromswold et al. 1996; Ben-Shachar et al. 2003; Bornkessel et al. 2005; Hagoort 2005; Friederici 2018). More recent proposals highlight the role of the LIFL in supporting retrieval from short term memory (Fiebach et al. 2005; Novick et al. 2005; Caplan et al. 2008; Leiken & Pylkkänen 2014). Moreover, LIFL is also implicating in studies on morphological processing (Sahin et al. 2006; Marslen-Wilson & Tyler 2007; Regel et al. 2017). Thus, activity in LIFL may again reflect one or many of the correlated processes involved with processing a bare NP vs. a case-inflected NP.

LATL is a key brain region in semantic processing, and has been implicated as a ‘hub’ for integration of conceptual features (Mollo et al. 2017; see Pylkkänen 2019a, 2019b for review). LATL shows greater activity for processing sentences compared to lists (Stowe et al. 1998; Rogalsky et al. 2009) and composing adjective-noun combinations compared to non-compositional contexts (Bemis & Pylkkänen 2011; Li & Pylkkänen 2021). Many studies also implicate OF (or ventromedial prefrontal cortex, vmPFC) as a key area in processing of semantic composition (Bemis & Pylkkänen 2011; Price et al. 2015; Binder et al. 2009, 2011). Thus, it is perhaps unsurprising that MEG activity in these areas correlated with the processing of an argument NP may differ as a function of their morphosyntactic features, as these features indirectly may affect the semantic composition of the sentence.

One key to interpreting the left inferior frontal and the left anterior temporal cortex activity is that the same brain regions appear to index case information and predict verb features. In addition to supporting language processing generally, LIFL may specifically be a key functional region in predicting grammatical features (Matchin et al. 2017). Similarly, as reviewed in Section 1.1., LATL and LOF activity correlate with the predictiveness of adjectives (*stainless*). Thus, greater activity in these regions specifically is consistent with the hypothesis that agreement processing involves early commitment or prediction of the expected verb features.

We observed an interaction between case features in Hindi in the LIPL, but not in Nepali. Crucially, the sentences in our experiment are virtually identical in terms of their syntactic structure, and the case morphemes in both languages serve virtually identical semantic and syntactic functions in the two languages, apart from their relevance for agreement. Thus, the processes that are correlated with the different case morphemes are controlled for between the two languages, and thus the effects reported here are unlikely to be due to some feature of accusative case marking in either language. Thus, we argue that the unique effect observed in this brain region in Hindi likely reflects the specific processes of selecting an agreement controller. However, we reported conflicting directions of the interaction between the whole-brain and targeted ROI analyses. In both analyses, we observed greater activity for the Ergative Subject NP, Accusative Object NP sentences (NP*-ne* NP*-ko*; default agreement configurations) compared to the Ergative Subject NP, Bare Object NP sentences (NP-*ne* NP; object agreement configurations), with no difference between the two Nominative Subject NP conditions (subject agreement configurations). However, the clusters diverged with respect to whether the subject agreement configurations patterned more like the default agreement configurations or the object agreement configurations. For our purposes, we predicted that there would be an interaction between Subject and Object NP Case features, but we had no prediction about the directionality of this interaction.

The LIPL appears to have a wide range of functions (Seghier 2013). In language processing, the LIPL is usually thought to be key to understanding event representations (Binder et al. 2009, Binder & Desai 2011) or semantic and thematic relations (Lewis et al. 2015; Thompson et al. 2007; Thothathiri et al 2012, Williams et al. 2017), to processing semantic composition (Boylan et al. 2015; Zhang et al. 2022), or to making logical inferences (Xu et al. 2024). Other work suggests that the LIPL may play a key role in processing of relations across sentences in discourse. For instance, Kandylaki et al. (2016) found a reduction in LIPL activity upon remention of the subject NP of a passive sentence vs. an active sentence, since passive voice is predictive of remention of the subject NP’s referent.

However, although agreement is a helpful cue in identifying the relations between an event and its participants, it’s not clear how to relate this line of research to the selection of a verb agreement controller. Outside of language processing, bilateral inferior parietal activity corresponds to attention shift (Corbetta & Shulman 1998; Singh-Curry & Husain 2009; Seghier 2013), especially attending to task-relevant “bottom-up” features (Ciaramelli et al. 2008). The supramarginal gyrus and angular gyrus may serve as a ‘hub’ for integration of processes in the prefrontal regions and posterior parts of the language network (Seghier et al. 2013; Kim et al. 2022).

One possible interpretation of these findings is that agreement controller selection in object agreement constructions can be conceived of as requiring a shift in attention. In Hindi, the generalization is that the most structurally prominent, morphologically bare argument NP is the agreement controller. In SOV structures, this requires participants to first attend to the morphological properties of the subject, integrate it into a syntactic analysis, then attend to the features of the object NP. In subject-agreement configurations, attention devoted to the morphosyntactic features of the subject NP are the same as the (expected) features of the verb, and attention devoted to the subject NP’s features while processing the subject NP can be ‘carried forward’ as the predictions of the verb’s features. In the critical object-agreement configurations, the subject NP’s features must be suppressed, and the inflectional features of the object NP instead must be ‘stored’ as the expected features of the verb. Crucially, processing the lexical features of both arguments must occur in order to interpret the sentence, thus subject-agreement configurations involve a conflict between processing the object NP and maintaining the inflectional features of the subject NP in memory. This is mitigated in the object-agreement configurations, in which the inflectional features of the subject NP need not be maintained for argument-verb agreement. This could explain the reduction in activity observed in the LIPL. Alternatively, the increase in activity for the default agreement sentences may index a conflict between attending to the features of the subject and object NPs, and instead retrieving a ‘default agreement’ rule to encode a default, 3^rd^ person singular masculine verb specification regardless of the features of any argument NP in the sentence.

If this is the right approach, our findings appear to conflict with Bhatia & Dillon’s (2022) behavioral findings. Bhatia & Dillon used the agreement-attraction paradigm, in which acceptability and processing time are affected by the presence of feature-matching NPs that are not agreement controllers (e.g. \**the key to the <u>cabinets</u> are on the table*; *Wagers* et al. 2009).

They found that Hindi users’ perception of acceptability of agreement errors in a verb was only modulated by NPs that controlled agreement for other verbs. In other words, morphological features (e.g., ‘bare’) are not the relevant cue for retrieving an agreement controller from memory, but rather a morphosyntactically unmarked feature that is assigned to argument NPs according to the principles of Hindi grammar. However, their findings crucially depend on processes occurring during or after the verb, whereas we report on evoked responses at the preverbal object NP. It is possible that Hindi comprehenders may initially evaluate an NP as a ‘candidate’ agreement controller based purely on morphological features, yielding the competition between subject and object NP features that we consider here. Afterwards, they may reject an NP as a possible agreement controller after integration into the syntactic analysis, rendering it invisible to attraction later. We leave reconciling these findings to future work.

Finally, the interpretation we propose here rests crucially on the generalization that only bare NPs are eligible for agreement controlling, and that Hindi comprehenders rely on this morphological cue. Other Indo-Aryan languages with split ergativity do not follow this generalization. For instance, Gujarati exhibits a nearly-identical case system as Hindi and Nepali. Like Hindi, the verb agrees with the object in the simple past, and this is cued by the presence of an ergative case suffix on the subject. However, unlike Hindi, the simple past agrees with the object NP regardless of whether it is bare, i.e., an accusative case suffix on the object does not ‘block’ object-verb agreement (Mistry 1997). If our interpretation of the Hindi data is correct, then we would expect a different interaction of case features in Gujarati. However, we would not predict that the pattern of activity would be the same, since accusative case features do not have the same relevance for identifying agreement controllers as in Hindi.

## 5. Conclusion

The human brain is capable of learning and representing any natural human language. Moreover, left fronto-temporal and temporo-parietal language regions subserve language comprehension across a variety of languages and language users. Yet, different grammatical rules may require that comprehenders store, attend to, or predict different properties of words and phrases, processes which must be supported by the brain. Additionally, users of different languages appear to integrate different cues into their interpretation of a sentence, reflecting different properties of their language. What is the relation between the otherwise-uniform ‘language network’, parametric grammatical differences, and language-specific adaptations that comprehenders exhibit as a function of their specific languages? In general, this is a difficult question to answer, because cross-language comparisons often confound many factors, i.e., very few languages only differ along one ‘parameter’ of variation. Here, we present on two parallel MEG experiments in Hindi and Nepali. Both languages feature similar case systems. Both languages use an aspect-based split-ergative case system, in which sentences with perfective verbs typically require an ergative subject, and other verbs require a nominative or ‘bare’ subject. Both languages also use a differential object marking system, in which object NPs are assigned accusative case if they are animate or specific inanimate NPs but otherwise may be unmarked.

However, the argument-verb system in Hindi is sensitive to the distribution of case features. Verbs must agree with the structurally highest NP that is bare. This means that there is no single cue that comprehenders can exploit to identify the agreement controller across sentence types. By contrast, Nepali exhibits subject-verb agreement consistently. Thus, by manipulating the distribution of case markers in Hindi and Nepali, we can identify brain activity associated with processing object-agreement configurations in Hindi, a grammatical rule that is absent in Nepali. This enables us to more squarely relate grammatical description, language processing strategy, and the organization of the brain’s language network to better understand how the ‘universal’ does the ‘language-specific’.

We observed activity in left inferior frontal lobe, left anterior temporal lobe, and orbitofrontal cortex corresponding to the processing of object NP case in both languages. In Hindi, we also observed that activity in these regions correlated with the inflections features of the expected verb, which we did not observe in Nepali. Additionally, we found that Hindi comprehenders exhibited an interaction of subject NP and object NP case features in the left inferior parietal lobe, and we failed to find a similar pattern in Nepali comprehenders. We suggested that these differences between Hindi and Nepali demonstrate the selective deployment of attention systems to identify a verb agreement controller. Our findings validate the utility of carefully controlled, cross-language experiments in the cognitive neuroscience of language, and the importance of research in less-studied languages for psycholinguistic models in general.

## Acknowledgments

We’d like to thank Liina Pylkkänen for feedback and comments.

To be completed upon completion.

## References

Abadie, Peggy. 1974. Nepali as an ergative language. Linguistics of the Tibeto-Burman Area 1(1), 156–177. 10.32655/LTBA.1.1.06v

Adachi, Y., Shimogawara, M., Higuchi, M., Haruta, Y., Ochiai, M. (2001). Reduction of non-periodic environmental magnetic noise in MEG measurement by continuously adjusted least squares method. IEEE Transactions on Applied Superconductivity, 11(1), 669–672. 10.1109/77.919433

Aldridge, E. (2008). Generative approaches to ergativity. Language and Linguistics Compass 2(5), 966–965. 10.1111/j.1749-818X.2008.00075.x

Badecker, W., Kumiak, F. (2007). Morphology, agreement and working memory retrieval in sentence production: Evidence from gender and case in Slovak. Journal of Memory and Language, 56(1), 65–85. 10.1016/j.jml.2006.08.004

Barr, D.J., Levy, R., Scheepers, C., Tily, H.J. (2013). Random effects structure for confirmatory hypothesis testing: Keep it maximal. Journal of Memory and Language, 68(3), 255–278. 10.1016/j.jml.2012.11.001

Bates, D., Mächler, M., Bolker, B., Walker, S. (2015). Fitting linear mixed-effects models using *lme4*. Journal of Statistical Software, 67(1), 1–48. 10.18637/jss.v067.i01

Baumgaertner, A., Weller, C., Büchel, C. (2002). Event-related fMRI reveals cortical sites involved in contextual sentence integration. Neuroimage, 16, 736–745. 10.1006/nimg.2002.1134

Bemis, D.K., Pylkkänen, L. (2011). Simple composition: A magnetoencephalography investigation into the comprehension of minimal linguistic phrases. Journal of Neuroscience, 31, 2801–2814. 10.1523/JNEUROSCI.5003-10.2011

Ben-Shachar, M., Hendler, T., Kahn, I., Ben-Bashat, D., Grodzinsky, Y. (2003). The neural reality of syntactic transformations: evidence from fMRI. Psychological Science, 14, 433–440. 10.1111/1467-9280.01459

Benjamini, Y., Hochberg, Y. (1995). Controlling the false discovery rate: a practical and powerful approach to multiple testing. Journal of the Royal Statistical Society: Series B (Methodological), 57(1), 289–300. 10.1111/j.2517-6161.1995.tb02031.x

Bhatia, S. (2019). Computing agreement in a mixed system. Ph.D. thesis, University of Massachusetts: Amherst. 10.7275/csbe-8c55

Bhatia, S., & Dillon, B.W. (2022). Processing agreement in Hindi: When agreement feeds attraction. Journal of Memory and Language, 125, 104322. 10.1016/j.jml.2022.104322

Bhatt, R. (2005). Long-distance agreement in Hindi-Urdu. Natural Language and Linguistic Theory, 23, 757–807. 10.1007/s11049-004-4136-0

Bhattamishra, S., Muralikrishnan, R., & Choudhary, K.K. (2021). Animacy modulates gender agreement comprehension in Hindi: An ERP study. *Language*, Cognition, and Neuroscience 37(5), 560–575. 10.1080/23273798.2021.1980219

Bickel, B. 2011. Grammatical relations typology. In J. J. Song (ed.), The Oxford Handbook of Language Typology. (pps. 399–445). Oxford: Oxford University Press.

Bickel, B., Witzlack-Makarevich, A., Choudhary, K. K., Schlesewsky, M., Bornkessel-Schlesewsky, I. (2015). The neurophysiology of language processing shapes the evolution of grammar: Evidence from case marking. PLoS ONE, 10(8), e0132819. 10.1371/journal.pone.0132819

Binder, J.R., Desai, R.H., Graves, W.W., Conant, L.L. (2009). Where is the semantic system? A critical review and meta-analysis of 120 functional neuroimaging studies. Cerebral Cortex, 19(12), 2767–2796. 10.1093/cercor/bhp055

Binder, J.R., Desai, R.H. (2011). The neurobiology of semantic memory. Trends in Cognitive Science, 15(11), 527–536. 10.1016/j.tics.2011.10.001

Bock, K., & Miller, C. A. (1991). Broken agreement. Cognitive Psychology, 23(1), 44–93. 10.1016/0010-0285(91)90003-7

Bornkessel, I., Zysset, S., Friederici, A.D., Yves von Cramon, D., Schlesewsky, M. (2005). Who did what to whom? The neural basis of argument hierarchies during language comprehension. NeuroImage, 26(1), 221–233. 10.1016/j.neuroimage.2005.01.032

Bornkessel-Schlesewsky, I., Schlesewsky, M. (2008). An alternative perspective on “semantic P600” effects in language comprehension. Brain Research Reviews 59(1), 55–73. 10.1016/j.brainresrev.2008.05.003

Bornkessel-Schlesewsky, I., Schlesewsky, M. (2009). The role of prominence information in the rela-time comprehension of transitive constructions: A cross-linguistic approach. Langauge and Linguistics Compass 59(1), 55–73. 10.1016/j.brainresrev.2008.05.003

Bornkessel-Schlesewsky, I., Krestzchmar, F., Tune, S., Wang, L., Genç, S., Philipp, M., Roehm, D., Schlesewsky, M. (2011). Think globally: Cross-linguistic variation in electrophysiological activity during sentence comprehension. Brain and Language 117(3), 133–152. 10.1016/j.bandl.2010.09.010

Bornkessel-Schlesewsky, I., Schlesewsky, M. (2016). The importance of linguistic typology for the neurobiology of language. Linguistic Typology 20(3), 615–621. 10.1515/lingty-2016-0032

Bornkessel-Schlesewsky, I., Schlesewsky, M. (2019). Toward a neurobiologically plausible model of language-related, negative event-related potentials. Frontiers in Psychology 21, 298. 10.3389/fpsyg.2019.00298

Boylan, C., Trueswell, J.C., Thompson-Schill, S.L. (2015). Compositionality and the angular gyrus: A multi-voxel similarity analysis of the semantic composition of nouns and verbs. Neuropsychologia 78, 130–141. 10.1016/j.neuropsychologia.2015.10.007

Brodbeck, C., Das, P., Singham, J., Reddigari, S., Brooks, T.L. (2023). Eelbrain 0.39. 10.5281/zenodo.7951251

Butt, M., & King, T.H. (2004). The status of case. In V. Dayal & A. Mahajan (eds.), Clause Structure in South Asian Languages. Studies in Natural Language and Linguistic Theory. (pp. 199–226). Dordrecht: Springer. 10.1007/978-1-4020-2719-2_6

Butt, M., & Poudel, T. 2007. Distribution of the ergative in Nepali. Manuscript, University of Kostanz.

Butt, M., & Deo, A. (2017). Developments into and out of ergativity: Indo-Aryan diachrony. In J. Coon, D. Massam, L. deMena Travis (eds.), The Oxford Handbook of Ergativity. (pp. 530–552). Oxford: Oxford University Pres. 10.1093/oxfordhb/9780198739371.013.22

Caplan, D., Stanczak, L., & Waters, G. (2008). Syntactic and thematic constraint effects on blood oxygenation level dependent signal correlates of comprehension of relative clauses. Journal of Cognitive Neuroscience, 20, 643–656. 10.1162/jocn.2008.20044

Carreiras, M., Duñabeitia, J.A., Vergara, M., de la Cruz-Pavia, I., Laka, I. (2010). Subject relative clauses are not universally easier to process: Evidence from Basque. Cognition 115(1), 79–92. 10.1016/j.cognition.2009.11.012

Carreiras, M., Carr, L., Barber, H.A., Hernández, A. (2010). Where syntax meets math: Right intraparietal sulcus activation in response to grammatical number agreement violations. NeuroImage, 49(2), 1741–1749. 10.1016/j.neuroimage.2009.09.058

Carreiras, M., Quiñones, I., Mancini, S., Hernández-Cabrera, J.A., & Barber, H. (2015). Verbal and nominal agreement: An fMRI study. NeuroImage, 120(15), 88–103. 10.1016/j.neuroimage.2015.06.075

Chacón, D. A. (2022). Default is different: Relations and representations in agreement processing. *Language*, Cognition, and Neuroscience, 37(6), 785–804. 10.1080/23273798.2021.2022172

Choudhary, K.K., Schlesewsky, M., Roehm, D., Bornkessel-Schlesewsky, I. (2009). The N400 as a correlate of interpretively relevant linguistic rules: Evidence from Hindi. Neuropsychologia, 47(13), 3012–3022. 10.1016/j.neuropsychologia.2009.05.009

Chow, W.-Y., Nevins, A., Carreiras, M. (2018). Effects of subject-case marking on agreement processing: ERP evidence from Basque. Cortex 99, 319–329. 10.1016/j.cortex.2017.12.009

Ciaramelli, E., Grady, C.L., Moscovitch, M. (2008). Top-dowm and bottom-up attention to memory: A hypothesis (AtoM) on the role of the posterior parietal cortex in memory retrieval. Neuropsychologia, 46(7), 1828–1851. 10.1016/j.neuropsychologia.2008.03.022

Corbetta, M., Shulman, G.L. (1998). Human cortical mechanisms of visual attention during orienting and search. Philosophical Transactions of the Royal Society B: Biological Sciences, 353(1373), 1353–1362. 10.1098/rstb.1998.0289

Cui, H., Jeong, H., Okamoto, K. Takahashi, D., Kawashima, R., & Sugiura, M. (2022). Neural correlates of Japanese honorific agreement processing mediated by socio-pragmatic factors: An fMRI study. Journal of Neurolinguistics, 62, Article 101041. 10.1016/j.jneuroling.2021.101041

Díaz, B., Erdocia, K., de Menezes, R.F., Mueller, J.L., Sebastián-Gallés, N., Laka, I. (2016). Electrophysiological correlates of second-language syntactic processes are related to native and second language distance regardless of age of acquisition. Frontiers in Psychology 7, 133. 10.3389/fpsyg.2016.00133

Dillon, B.W., Mishler, A., Sloggett, S., & Phillips, C. (2013). Contrasting intrusion profiles for agreement and anaphora: Experimental and modeling evidence. Journal of Memory and Language, 69(2), 85–103. 10.1016/j.jml.2013.04.003

Dixon, R.M.W. (1994). Ergativity. Cambridge: Cambridge University Press. 10.1017/CBO9780511611896

Dunagan, D., Zhang, S., Li, J., Bhattasali, S., Pallier, C., Whitman, J., Yang, Y., Hale, J. (2022). Neural correlates of semantic number: A cross-linguistic investigation. Brain and Language 229, 105110. 10.1016/j.bandl.2022.105110

Eberhard, K.M., Cutting, J.C., Bock, K. (2005). Making sense of syntax: Number agreement in sentence production. Psychological Review, 112(3), 531–559. 10.1037/0033-295X.112.3.531

Ferreira, F., Qiu, Z. (2021). Predicting syntactic structure. Brain Research 1770, 147632. 10.1016/j.brainres.2021.147632

Fiebach, C.J., Schlesewsky, M., & Friederici, A.D. (2002). Separating syntactic memory costs and syntactic integration costs during parsing: The processing of German WH-questions. Journal of Memory and Language, 47(2), 250–272. 10.1016/S0749-596X(02)00004-9

Franck, J., Vigliocco, G., Nicol, J. (2002). Subject-verb agreement errors in French and English: The role of syntactic hierarchy. Language and Cognitive Processes, 17(4), 371–404. 10.1080/01690960143000254

Friederici, A. (2002). Towards a neural basis of auditory sentence processing. Trends in Cognitive Science, 6, 78–84. 10.1016/S1364-6613(00)01839-8

Friederici, A. (2018). The neural basis for human syntax: Broca’s area and beyond. Current Opinion in Behavioral Sciences, 21, 88–92. 10.1016/j.cobeha.2018.03.004

Fruchter, J., Linzen, T., Westerlund, M., Marantz, A. (2015). Lexical preactivation in basic linguistic phrases. Journal of Cognitive Neuroscience 27(10), 1912–1935. 10.1162/jocn_a_00822

Gramfort, A., Luessi, M., Larson, E., Engemann, D.A., Strohmeier, D., Brodbeck, C., Goj, R., Jas, M., Brooks, T., Parkkonen, L., Hämäläinen, M.S. (2013). MEG and EEG data analysis with MNE-Python. Frontiers in Neuroscience, 7(267), 1–13. 10.3389/fnins.2013.00267

Gulati, M., Choudhary, K. K. (2023). Cross-linguistic variations in the processing of ergative case: Evidence from Punjabi. In P. Chandra (eds), Variation in South Asian Languages, 267–294. Dordrecht: Springer. 10.1007/978-981-99-1149-3

Gulati, M., Muralikrishnan, R., Choudhary, K.K. (2024). An ERP study on the processing of subject-verb and object-verb gender agreement in Punjabi. Journal of Psycholinguistic Research 53(4), 59. 10.1007/s10936-024-10095-4

Hagoort, P. (2005). On Broca, brain, and binding: A new framework. Trends in Cognitive Sciences, 9(9), 416–423. 10.1016/j.tics.2005.07.004

Hagoort, P., Brown, C., & Groothusen, J. (1993). The syntactic positive shift (SPS) as an ERP measure of syntactic processing. Language and Cognitive Processes, 8, 439–483. 10.1080/01690969308407585

Halgren, E, Dhond, R.P., Christensen, N., Van Petten, C., Marinkovic, K., Lewine, J.D., Dale, A.M. (2002). N400-like magnetoencephalography responses modulated by semantic context, word frequency, and lexical class in sentences. NeuroImage, 17(3), 1101–1106. 10.1006/nimg.2002.1268

Hauptman, M., Blanco-Elorrieta, E., & Pylkkänen, L. (2021). Inflection across categories: Tracking abstract morphological processing in language production with MEG. Cerebral Cortex, 2021, Article bhab309. 10.1093/cercor/bhab309

de Hoop, H., Narasimhan, B. (2009). Ergative Case-marking in Hindi. In H. de Hoop & P. de Swart (eds.), Differential Subject Marking, 63–78. Dordrecht: Springer. 10.1007/978-1-4020-6497-5_4

Kachru, Y. 1980. Aspects of Hindi Grammar. New Delhi: Manohar Publications.

Kandylaki, K.D., Nagels, A., Tune, S., Kircher, T., Wiese, R., Schlesewsky, M., Bornkessel-Schlesewsky, I. (2016). Predicting ‘when’ in discourse engages the human dorsal auditory system: An fMRI study using naturalistic stories. Journal of Neuroscience 36(48), 12180–12191. 10.1523/JNEUROSCI.4100-15.2016

Keshev, M., Meltzer-Asscher, A. (2024). The representation of agreement features in memory is updated during sentence processing: Evidence for verb-reflexive interactions. Journal of Memory and Language, 135, 104495. 10.1016/j.jml.2023.104495

Kielar, A., Milman, L., & Bonakdarpour, B. (2011). Neural correlates of covert and overt production of tense and agreement morphology: Evidence from fMRI. Journal of Neurolinguistics, 24(2), 183–201. 10.1016/j.jneuroling.2010.02.008

Kim, H., Wang, K., Cutting, L.E., Willcut, E.G., Petrill, S.A., Leopold, D.R., Reineberg, A.E., Thompson, L.A., Banich, M.T. (2022). The angular gyrus as a hub for modulation of language-related cortex by distinct prefrontal executive control regions. Journal of Cognitive Neuroscience 34(12), 2275–2296. 10.1162/jocn_a_01915

Kriegeskorte, N., Mur, M., Bandettini, P. (2008). Representational similarity analysis – connecting the branches of systems neuroscience. Frontiers in Systems Neuroscience 2, 4. 10.3389/neuro.06.004.2008

Kuroda, S.-Y. (1972). The categorical and the thetic judgment: Evidence from Japanese syntax. Foundations of Language 9(2), 153–185.

Kutas, M., & Hilyard, S. A. (1980). Reading senseless sentences: Brain potentials reflect semantic incongruity. Science, 207(4427), 203–205. 10.1126/science.735065

Kutas, M., & Federmeier, K. (2011). Thirty years and counting: Finding meaning in the N400 component of the event-related brain potential (ERP). Annual Review of Psychology, 62, 621– 647. 10.1146/annurev.psych.093008.131123

Laka, I. (2012). Merging from the temporal input: On subject-object asymmetries and an ergative language. In M. Piattelli-Palmerini & R.C. Berwick (eds.), Rich Languages from Poor Inputs, 127–145. Oxford: Oxford University Press. 10.1093/acprof:oso/9780199590339.003.0009.

Lau, E., Phillips, C., & Poeppel, D. (2008). A cortical network for semantics: (De)constructing the N400. Nature Reviews: Neuroscience, 9, 920–933. 10.1038/nrn2532

Lau, E., & Namyst, A. (2019). fMRI evidence that left posterior temporal cortex contributes to N400 effects of predictability independent of congruity. Brain and Language, 199, Article 104697. 10.1016/j.bandl.2019.104697

Leiken, K., Pylkkänen, L. (2014). MEG evidence that the LIFG effect of object extraction requires similarity-based interference. *Language*, Cognition, and Neuroscience, 29(3), 381–389. 10.1080/01690965.2013.863369

Lewis, G.A., Poeppel, D., Murphy, G.L. (2015). The neural bases of taxonomic and thematic conceptual relations: An MEG study. Neuropsychologia, 68, 176–189. 10.1016/j.neuropsychologia.2015.01.011

Li, C. (2007). Split ergativity and split intransitivity in Nepali. Lingua, 117, 1462–1482. 10.1016/j.lingua.2006.09.002

Li, J., Pylkkänen, L. (2021). Disentangling semantic composition and semantic association in the left temporal lobe. Journal of Neuroscience 41(30), 6526–6538. 10.1523/JNEUROSCI.2317-20.2021

Lindemann, L. 2019. A jewel inlaid: Ergativity and markedness in Nepali. PhD Thesis, Yale University.

Logenbaugh, N., Polinsky, M. (2016). The processing of long-distance dependencies in Niuean. In The Proceedings of Austronesian Formal Linguistics Association (AFLA) 22. https://hdl.handle.net/1885/101156

Mahajan, A. (2012). Ergatives, antipassives, and the overt light v in Hindi. Lingua 122(3), 204–214. 10.1016/j.lingua.2011.10.011

Mahajan, A. (2017). Accusative and ergative in Hindi. In J. Coon (ed.), The Oxford Handbook of Ergativity, 86–108. Cambridge: Oxford University Press. 10.1093/oxfordhb/9780198739371.013.4

Malik-Moraleda, S., Ayyash, D., Gallée, J., Affourtit, J., Hoffmann, M., Mineroff, Z., Jouravlev, O., Fedorenko, E. (2022). An investigation across 45 languages and 12 language families reveals a universal language network. Nature: Neuroscience, 25, 1014–1019. h#ps://doi.org/10.1038/s41593-022-01114-5

Marslen-Wilson, W. D., Tyler, L. K. (2007). Morphology, language, and the brain: The decompositional substrate for language comprehension. Philosophical Transactions of the Royal Society: Biological Sciences, 362(1481), 823–836. 10.1098/rstb.2007.2091

Matchin, W., Hammerly, C., Lau, E. (2017). The role of the IFG and pSTS in syntactic prediction: Evidence from a parametric study of hierarchical structure in fMRI. Cortex 88, 106–123. 10.1016/j.cortex.2016.12.010

Matthew, A.M., Muralikrishnan, R., Gulati, M., Bhattamishra, S., Choudhary, K.K. 2024. Processing subject and object agreement in light verb constructions: Electrophysiological insights from Hindi. Poster presented at the Human Sentence Processing Conference. Ann Arbor, MI.

Mistry, P. (1997). Objecthood and specificity in Gujarati. In L. Campbell, J. Hill, & P.J. Mistry (eds.), The Life of Language: Papers in Honor of William Bright. Berlin: Mouton de Gruyter. 10.1515/9783110811155.425

Mollo, G., Cornellisen, P.L., Millman, R.E., Ellis, A.W., Jeffries, E. (2017). Oscillatory dynamics supporting semantic cognition: MEG evidence for the contribution of the anterior temporal lobe hub and modality-specific spokes. PLoS ONE 12, e0169269. 10.1371/journal.pone.0169269

Moitra, S., Chacón, D.A., Stockall, L. (2024). How long is long?: Word length effects in reading correspond to minimal graphemic units: An MEG study in Bangla. PLoS ONE 19(4), e0292979. 10.1371/journal.pone.0292979

Molinaro, N., Barber, H.A., Carreiras, M. (2011). Grammatical agreement processing in reading: ERP findings and future directions. Cortex, 47(8), 908–930. 10.1016/j.cortex.2011.02.019

Nieuwland, M.S., Martin, A.E., & Carreiras, M. (2011). Brain regions that process case: Evidence from Basque. Human Brain Mapping, 33(11), 2509–2520. 10.1002/hbm.21377

Nevins, A., Dillon, B., Malhotra, S., & Phillips, C. (2007). The role of feature-number and feature-type in processing Hindi verb agreement violations. Brain Research, 1164, 81–94. 10.1016/j.brainres.2007.05.058

Nicol, J.L., Forster, K.I., & Veres, C. (1997). Subject-verb agreement process in comprehension. Journal of Memory and Language, 36(4), 569–587. 10.1006/jmla.1996.2497

Novick, J.M., Trueswell, J.C., & Thompson-Schill, S.L. (2005). Cognitive control and parsing: Reexamining the role of Broca’s area in sentence comprehension. Cognitive, Affective & Behavioral Neuroscience, 5, 263–281. 10.3758/CABN.5.3.263

Pandharipande, R., & Kachru, Y. (1977). Relational grammar, ergativity, and Hindi-Urdu. Lingua, 41, 217–238. 10.1016/0024-3841(77)90080-8

Philipp, M., Bornkessel-Schlesewsky, I., Bisang, W., Schlesewsky, M. (2008). The role of animacy in the real time comprehension of Mandarin Chinese: Evidence from auditory event-related potentials. Brain and Language 105(2), 112–133. 10.1016/j.bandl.2007.09.005

Polinsky, M., Gallo, C.G., Graff, P., Kravtchenko, E. (2012). Subject preference and ergativity. Lingua 122(3), 267–277. 10.1016/j.lingua.2011.11.004

Price, A., Bonner, M.F., Peelle, J.E., Grossman, M. (2015). Converging evidence for the neuroanatomic basis of combinatorial semantics in the angular gyrus. Journal of Neuroscience 35, 3276–3284. 10.1523/JNEUROSCI.3446-14.2015

Pylkkänen, L., Llinás, R., McElree, B. (2004). Distinct effects of semantic plausibility and semantic composition in MEG. In Proceedings of the 14th International Conference on Biomagnetism. Boston, MA: Biomag.

Pylkkänen, L. (2019a). The neural basis of combinatory syntax and semantics. Science, 366(6461), 62–66. 10.1126/science.aax0050

Pylkkänen, L. (2019b). Neural basis of basic composition: What we have learned from the red-boat studies and their extensions. Philosophical Transactions of the Royal Society B 375, 20190299. 10.1098/rstb.2019.0299

Quiñones, I., Molinaro, N., Mancini, S., Hernández-Cabrera, J.A., Carreiras, M. (2014). Where agreement merges with disagreement: fMRI evidence of subject-verb integration. NeuroImage 88, 188–201. 10.1016/j.neuroimage.2013.11.038

Quiñones, I., Molinaro, N., Mancini, S., Hernández-Cabrera, J.A., Barber, H., Carreiras, M. (2018). Tracing the interplay between syntactic and lexical features: fMRI evidence from agreement comprehension. NeuroImage 175, 259–271. 10.1016/j.neuroimage.2018.03.069

R Core Team, 2021. R: A language and environment for statistical computing. R Foundation for Statistical Computing, Vienna, Austria. https://www.R-project.org

Regel, S., Kotz, S. A., Henseler, I., Friederici, A.D. (2017). Left inferior frontal gyrus mediates morphosyntax: ERP evidence from verb processing in left-hemisphere damaged patients. Cortex, 86, 156–171. 10.1016/j.cortex.2016.11.007

Rogalsky, C., Saberi, K., Hickok, G. (2009). Temporal and structural contributions to activation of anterior temporal sentence processing regions: an fMRI study. Structure 55, 9. 10.1016/s1053-8119(09)71759-8

Sahin, N.T., Pinker, S., Halgren, E. (2006). Abstract grammatical processing of nouns and verbs in Broca’s area: Evidence from fMRI. Cortex, 42, 540–562. 10.1016/S0010-9452(08)70394-0

Sahin, N.T., Pinker, S., Cash, S.S., Schomer, D., Halgren, E. (2009). Sequential processing of lexical, grammatical, and phonological information within Broca’s Area. Science, 326(5951), 445–459. 10.1126/science.1174481

Sassenhagen, J., & Draschkow, D. (2019). Cluster-based permutation tests of MEG/EEG data do not establish significance of effect latency or location. Psychophysiology, 56(6), e13335. 10.1111/psyp.13335

Schelewesky, M., Bornkessel-Schlesewsky, I. (2009). When semantic P600s turn into N400s: On cross-linguistic differences in online verb-argument linking. In K. Alter, M. Horne, M. Lindgren, M. Roll, J. von Koss Torkildsen (eds.), Proceedings of Brain Talk, 75–100. Lund: Lunds Universitet.

Seghier, M.L. (2013). The angular gyrus: Multiple functions and multiple subdivisions. The Neuroscientist, 19(1), 43–61. 10.1177/107385841244059

Simonsen, R., Chacón, D.A. (2024). Using word order cues to predict verb class in L2 Spanish. Bilingualism: Language and Cognition. To appear.

Singh-Curry, V., Husan, M. (2009). The functional role of the inferior parietal lobe in the dorsal and ventral stream dichotomy. Neuropsychologia, 47(6), 1434–1448. 10.1016/j.neuropsychologia.2008.11.033

Slade, B. 2014. The diachrony of light and auxiliary verbs in Indo-Aryan. Diachronica 30(4), 531–578. 10.1075/dia.30.4.04sla

Solomyak, O., Marantz, A. (2010). Evidence for early morphological decomposition in visual word recognition. Journal of Cognitive Neuroscience 22(9), 2042–2057. 10.1162/jocn.2009.21296

Stromswold, K., Caplan, D., Alpert, N., Rauch, S. (1996). Localization of syntactic comprehension by positron emission tomography. Brian and Language, 52, 452–473. 10.1006/brln.1996.0024

Stowe, L.A., Broere, C.A., Paans, A.M., Wijers, A.A., Mulder, G., Vaalburg, W., Zwarts, F. (1998). Localizing components of a complex task: Sentence processing and working memory. Neuroreport 9, 2995–2999. 10.1097/00001756-199809140-00014

Thompson, C.K., Bonakdarpour, B., Fix, S.C., Blumenfeld, H.K., Parrish, T.B., Gitelman, D.R., Mesulam, M.M. (2007). Neural correlates of verb argument structure processing. Journal of Cognitive Neuroscience, 19(11), 1753–1767. 10.1162/jocn.2007.19.11.1753

Thothathiri, M., Kimberg, D.Y., Schwartz, M.F. (2012). The neural basis of reversible sentence comprehension: Evidence from voxel-based lesion symptom mapping in aphasia. Journal of Cognitive Neuroscience, 24(1), 212–22. 10.1162/jocn_a_00118

Tollan, R., Massam, D., Heller, D. (2019). Effects of case and transitivity on processing dependencies: Evidence from Nieuan. Cognitive Science 43(6), e12736. 10.1111/cogs.12736

Tucker, M., Politzer-Ahles, S., King, J., & Almeida, D. (2014). Agreement attraction in the neural language system. Poster presented at AMLaP 20, Edinburgh.

Van Petten, C., & Luka, B.J. (2006). Neural localization of semantic context effects in electromagnetic and hemodynamic studies. Brain and Language, 97(3), 279–293. 10.1016/j.bandl.2005.11.003 https://doi.org/10.1016/j.bandl.2005.11.003

Williams, A., Reddigari, S., & Pylkkänen, L. (2017). Early sensitivity of left perisylvan cortex to relationality in nouns and verbs. Neuropsychologia, 100, 131–143. 10.1016/j.neuropsychologia.2017.04.029

Wagers, M.W., Lau, E., & Phillips, C. (2009). Agreement attraction in comprehension: Representations and processes. Journal of Memory and Language, 61(2), 206–237. 10.1016/j.jml.2009.04.002

Wang, L., Schlesewsky, M., Bickel, B., Bornkessel-Schlesewsky, I. (2009). Exploring the nature of the subject preference: Evidence from the online comprehension of simple sentences in Mandarin Chinese. Language and Cognitive Processes 24(7–8), 1180–1126.

Wang, L., Schlesewsky, M., Philipp, M., Bornkessel-Schlesewsky, I. (2012). The role of animacy in online argument interpretation in Mandarin Chinese. In M. Lamers & P. de Swart (eds.), Case, Word Order, and Prominence, 91–119. Netherlands: Springer. 10.1007/978-94-007-1463-2_5

Wei, X., Adamson, H., Schwendemann, M., Goucha, T., Friederici, A.D., Anwander, A. (2023). Native language differences in the structural connectome of the human brain. NeuroImage 270, 119955. 10.1016/j.neuroimage.2023.119955

Xu, B., Wu, J., Xiao, H., Fünte, T.M., Ye, Z. (2024). Inferior parietal cortex represents relational structures for explicit transitive inference. Cerebral Cortex 34(4), bhae137. 10.1093/cercor/bhae137

Yokoyama, S., Maki, H., Hashimoto, Y., Toma, M., Kawashima, R. (2012). Mechanism of case processing in the brain: An fMRI study. PLoS ONE 7(7), e40474. 10.1371/journal.pone.0040474

Zawizsewski, A., Friederici, A.D. (2009). Processing canonical and non-canonical sentences in Basque: The case of object-verb agreement as revealed by event-related brain potentials. Brain Research 1284, 161–179. 10.1016/j.brainres.2009.05.099

Zhang, W., Xiang, M., Wang, S. (2022). The role of the left angular gyrus in the representation of linguistic composition rules. Human Brain Mapping 43(7), 2204–2217. 10.1002/hbm.25781

